# Perceptual gating of a brainstem reflex facilitates speech understanding in human listeners

**DOI:** 10.1101/2020.05.31.115444

**Authors:** Heivet Hernandez-Perez, Jason Mikiel-Hunter, David McAlpine, Sumitrajit Dhar, Sriram Boothalingam, Jessica J.M. Monaghan, Catherine M. McMahon

## Abstract

Navigating “cocktail party” situations by enhancing foreground sounds over irrelevant background information is typically considered from a cortico-centric perspective. However, subcortical circuits, such as the medial olivocochlear (MOC) reflex that modulates inner ear activity itself, have ample opportunity to extract salient features from the auditory scene prior to any cortical processing. To understand the contribution of auditory subcortical nuclei and the cochlea, physiological recordings were made along the auditory pathway while listeners differentiated non(sense)-words and words. Both naturally-spoken and intrinsically-noisy, vocoded speech — filtering that mimics processing by a cochlear implant—significantly activated the MOC reflex, whereas listening to speech-in-background noise revealed instead engagement of midbrain and cortical resources. An auditory periphery model reproduced these speech degradation-specific effects, providing a rationale for goal-directed gating of the MOC reflex to enhance representation of speech features in the auditory nerve. Our data reveals the co-existence of two strategies in the auditory system that may facilitate speech understanding in situations where the speech signal is either intrinsically degraded or masked by extrinsic auditory information.

## Introduction

Cocktail-party listening, the ability to focus on a single talker in a background of simultaneous, overlapping conversations, is critical to human communication, and a long-sought goal of hearing technologies [1, 2]. Problems listening in background noise are a key complaint of many listeners with even mild hearing loss, and a stated factor in the non-use and non-uptake of hearing devices [3–5]. However, despite its importance in every-day listening tasks, and its relevance to hearing impairment, physiological mechanisms that enhance attended speech remain poorly understood. In addition to local circuits in the auditory periphery and brainstem that have evolved to enhance automatically the neural representation of ecologically relevant sounds [6–8], it is likely that such a critical goal-directed behaviour as cocktail-party listening also relies on top-down, cortically-driven processes to emphasize perceptually relevant sounds, and suppress those that are irrelevant [9, 10]. Nevertheless, the specific role of bottom up and top-down mechanisms in complex listening tasks remain to be determined.

A potential mechanistic pathway supporting cocktail party listening is the auditory efferent system, whose multi-synaptic connections extend from auditory cortex (AC) to the inner ear [11–13]. In particular, reflexive activation of fibers in the medial olivocochlear (MOC) reflex innervating the outer hair cells (OHCs)—electromotile elements responsible for the cochlea’s active amplifier—is known to reduce cochlear gain [14], thereby increasing the overall dynamic range of the inner ear and facilitating sound encoding in high levels of background noise [15].

MOC fibers (ipsilateral and contralateral to each ear) originate in medial divisions of the superior olivary complex in the brainstem and synapse directly on the basal aspects of OHCs (Warr and hand neural sensitivity to sound [16, 17].

MOC neurons are also innervated by descending fibers from AC and midbrain neurons, providing potential means of modulating cochlear activity in a task-related or attention-dependent manner [18–20]. Although it has been speculated that MOC-mediated changes in cochlear gain might enhance speech coding in background noise [21], the role of the MOC reflex in reducing cochlear gain during goal-directed listening in normal hearing human subjects (i.e., with physiologically normal OHCs) remains controversial. In particular, it remains unclear under which conditions MOC reflex is active, including whether listeners must actively be engaged in listening task [22–24].

Efferent-mediated changes in cochlear gain can be assessed by measuring otoacoustic emissions (OAEs), energy generated by the active OHCs and measured non-invasively as sound from the ear canal [25]. When transient sounds, such as clicks, are delivered to one ear in the presence of noise in the opposite ear; OAE amplitudes are reduced, reflecting increased MOC efferent activity [26]. The magnitude of OAE s has been reported as either increased [23, 27], reduced [22, 28] or unaffected [29, 30] in participants with improved speech-in-noise perception. Confounding effects on cochlear gain could depend on factors such as task difficulty or relevance (e.g., speech *vs.* non-speech tasks) or even methodological differences in recording and analysing inner ear signatures such as OAEs [23, 24].

Here, we examined the role of the auditory efferent system in active (participant’s attention directed towards the speech stimuli) *vs*. passive (participant’s attention directed away from the speech stimuli and towards a silent, non-subtitled film) listening conditions for three commonly employed speech manipulations: vocoding of ‘natural’ speech— filtering that mimics processing by a cochlear implant; speech presented in a background of multi-talker ‘babble’ noise, and speech presented in a background of speech-shaped noise i.e. noise with the same long-term spectrum as speech. Physiological recordings in the central auditory pathway, including brainstem, midbrain and cortical responses were made whilst listeners performed an active listening task (detecting non-words in a string of Australian-English words and non-words). Our experimental paradigm was designed to maintain fixed levels of task difficulty that allowed us to preserve comparable task relevance across the speech manipulations and therefore avoid confounding effects of task difficulty on top-down modulation of activation of the MOC reflex. In addition, homologous visual and auditory scenes were implemented to control for differences in alertness between active and passive listening conditions.

When task difficulty was maintained across speech manipulations, measures of hearing function at the level of the cochlea, brainstem, midbrain and cortex were modulated differentially depending on the type of degradation applied to speech sounds, and on whether or not speech was actively attended. Specifically, the MOC reflex, assessed in terms of the magnitude of click-evoked OAEs (CEOAEs), was activated by vocoded speech—an intrinsically degraded speech signal—but not by otherwise ‘natural’ speech presented in either babble-noise or speech-shaped noise. Furthermore, neural activity generated by the auditory midbrain was significantly increased in active *vs*. passive listening for speech in babble and speech-shaped noise, but not for vocoded speech. This increase was associated with elevated cortical markers of listening effort for the speech-in-noise conditions. A model of the auditory periphery including an MOC circuit with biophysically realistic temporal dynamics confirmed the stimulus-dependent role of the MOC reflex in enhancing neural coding of speech signals. Our data suggest that otherwise identical performance in active listening tasks may invoke quite different efferent circuits, requiring different levels of listening effort, depending on the type of stimulus degradation experienced.

## Results

### Maintaining task relevance across speech manipulations requires iso-performance

We assessed speech intelligibility—specifically the ability to discriminate between Australian-English words and non-words—when speech was degraded by three different manipulations: noise-vocoding the entire speech waveform; adding 8-talker babble noise to ‘natural’ speech (i.e. speech-in-quiet) and adding speech-shaped noise to ‘natural’ speech. Participants were asked to respond (by means of a button press) when they heard a non-word in a string of words and non-words (Figure1A).

Three levels of task difficulty were achieved by altering either the number of noise-vocoded channels—16 (Voc16), 12 (Voc12) and 8 (Voc8) channels)—or by altering the signal-to-noise ratio (SNR) when speech was masked by babble noise—+10 (BN10) and +5 (BN5) dB SNR—or speech-shaped noise—+8 (SSN8) and +3 (SSN3) dB SNR (Figure 1B). This was statistically confirmed in all 56 listeners (n=27 in the vocoded condition, and n=29 in the two masked conditions) who showed consistently better performance—a higher rate of detecting non-words—in the less degraded conditions i.e. more vocoded channels or higher SNRs in the masked manipulations (Repeated Measures ANOVA (rANOVA) [vocoded: [F (3, 78) = 70.92, p = 0.0001]; babble noise: [F (3, 78) = 70.92, p = 0.0001] and speech-shaped noise: [F (2, 56) = 86.23, p = 0.0001], *post hoc* analysis with bonferroni corrections (6 multiple comparisons for the noise-vocoded experiment and 3 for BN and SSN manipulations), Figure 1B.

**Figure 1.**
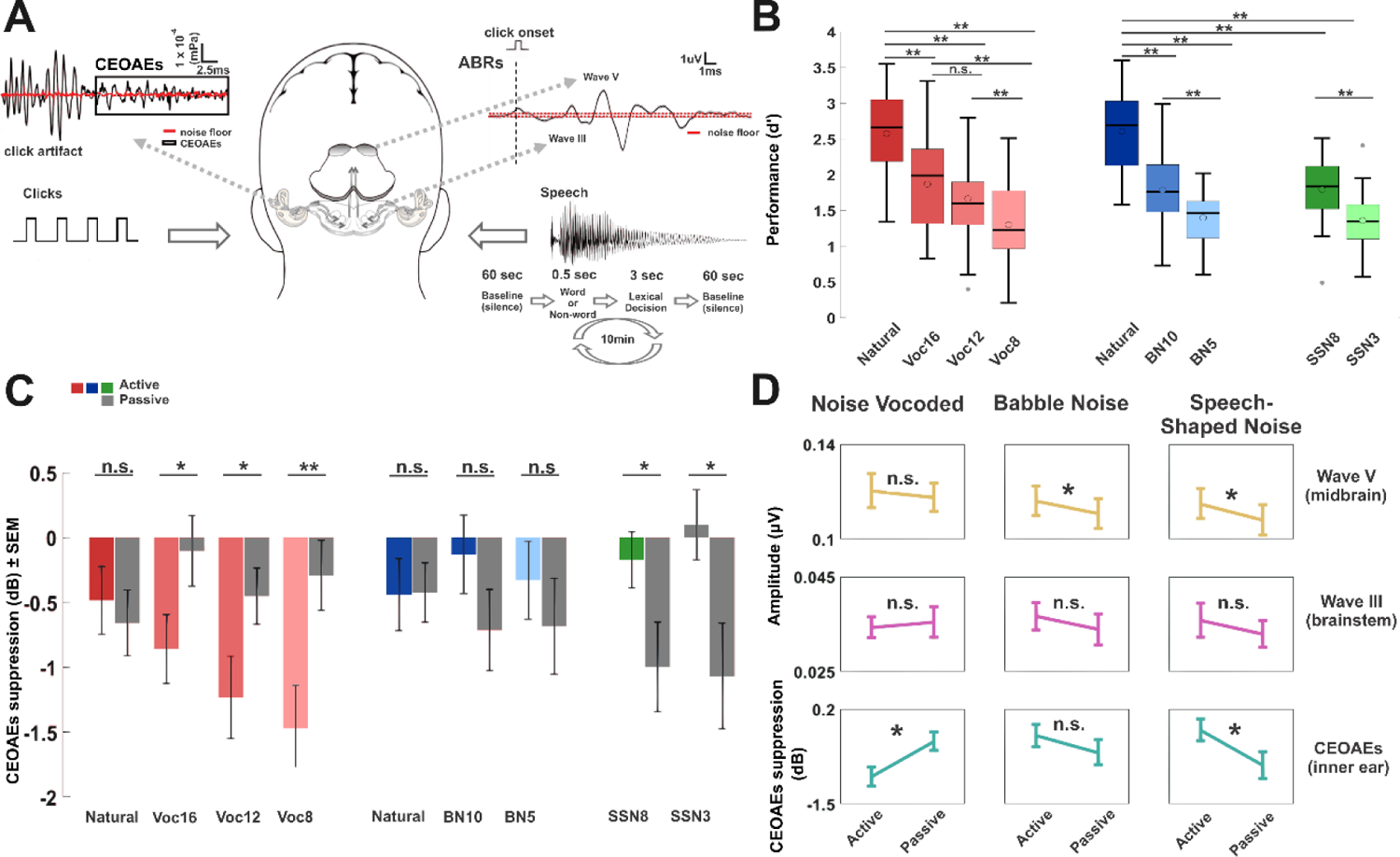
Behavioural and physiological measurements during active and passive listening. **A.** Schematic of the experimental paradigm (adapted with permissions, [31]). Clicks were continuously presented in one ear whereas speech tokens were delivered contralaterally in the other ear for 10 minutes to evoke CEOAEs. One minute of baseline CEOAEs (only clicks being delivered) were always recorded at the beginning and at the end of each condition. Participants had 3 sec to make a lexical-decision (discriminate words *vs.* non-words) in the active listening condition whereas in the passive condition they were asked to ignore all the auditory stimuli and watch a movie. Upper left panel shows the CEOAEs analysis window obtained from the inner ear recordings. Upper right panel corresponds to click-evoked ABRs recordings. **B**. Performance during the lexical-decision task. Mean d’ is represented in white circles (n=27 in the noise-vocoded condition, and n=29 in the two masked conditions), median corresponds to the horizontal line. The upper and lower limits of the boxplot represent the 1^st^ (q1) and 3^rd^ (q3) quartiles respectively while the whiskers denote the interquartile range (IQR =q3-q1). Within speech conditions comparisons showed that the highest performance was always achieved for natural speech compared to the all noise-vocoded, babble noise and speech-shaped noise manipulations *(bonferroni corrected* p *=* 0.001). Performance was moderately-high (statistically lower than natural speech but higher than Voc8, BN5 and SSN3) for Voc16 and Voc12 (non-significant differences (n.s.), *bonferroni corrected* p = 0.11) as well as for BN10 and SSN8 respectively. The lowest level of performance was predictably observed for Voc8, BN5 and SSN3 (p =0.001). **C.** Click-Evoked OAEs suppression. The figure shows mean CEOAE magnitude changes relative to the baseline for all conditions. **D.** CEOAEs and ABRs collapsed (rANOVA main effect of Conditions (Active and Passive)). CEOAEs and ABRs are reported as means ± SEM. (\*\**bonferroni corrected* p < 0.01; * *bonferroni corrected* p < 0.05).

Iso-performance was achieved across speech manipulations, with best performance observed in the two natural speech conditions—one employed in the vocoding experiment and the other in the two masked conditions: (One-way ANOVA [F (1, 54) = 0.43, p *=* 0.84]). A moderate and similar level of performance (significantly lower than performance for natural speech) was achieved across Voc16/ Voc12 (n.s. compared to each other; p *=* 0.11), BN10 and SSN8 conditions: (One-way ANOVA: [F (3, 108) = 0.67, p= 0.57]). The poorest performance, significantly lower than the high and moderate performance levels, was observed for Voc8, BN5 and SSN3: ([F (3, 81) = 0.07, p = 0.72]).

As increasing task difficulty has been previously linked to the allocation of auditory attention and cognitive resources towards the task itself [32], we employed the discrete and matching levels of task difficulty across speech manipulations as a proxy for the required auditory attention.

### MOC reflex is modulated by task engagement in a stimulus-dependent manner

To determine whether auditory attention modulates cochlear gain via the auditory efferent system in a task-dependent manner, we assessed the effect of active *vs*. passive listening and speech manipulation on inner ear activity. Click-evoked OAEs (CEOAEs) were recorded continuously whilst participants both actively performed the lexical task, and passively listened to the same *corpus* of speech tokens (Figure 1A). Compared to baseline measures obtained in the absence of speech (in the ipsilateral ear but click stimuli in the contralateral one), CEOAEs were significantly reduced in magnitude when actively listening to natural speech, and in all noise-vocoded conditions (natural: [t (24) = −2.33, p = 0.03]; Voc16: [t (23) = 3.40, p = 0.002]; Voc12: [t (24) = 3.98, p = 0.001] and Voc8: [t (25) = 5.14, p = 0.001]). Conversely, during passive listening, CEOAEs obtained during natural, but not noise-vocoded, speech were significantly smaller than baseline: [t (25) = 2.29, p = 0.03], as were CEOAEs recorded during the two masked conditions at all SNRs (natural: [t (26) = 2.17, p = 0.04]; BN10: [t (28) = 2.80, p = 0.009] and BN5: [t (28) = 2.36, p = 0.02]; SSN8: [t (28) = 3.37, p = 0.002] and SS3: [t (28) = 3.50, p = 0.002]). This suggests that auditory efferent activity is modulated differently in active and passive listening, and by the different speech manipulation types, despite iso-performance across experiments.

We calculated the reduction in CEOAEs between baseline and experimental conditions (CEOAE suppression)—a proxy for activation of the MOC reflex—to quantify auditory efferent control of cochlear gain in active and passive listening. For noise-vocoded speech, suppression of CEOAEs was significantly greater when participants were actively engaged in the lexical task (−1.01 ± 0.18 dB) compared to when they were asked to ignore the auditory stimuli (−0.38 ± 0.18 dB): rANOVA: [F (1, 22) = 8.49, p = 0.008], (Figure 2A). Moreover, a significant interaction was observed between conditions and stimulus-type: [F (3, 66) = 2.80, p = 0.046], indicating that the suppression of CEOAEs was stronger for all vocoded conditions in which listeners where required to make lexical decisions, compared to when they were not—Voc16: ([t (23) = −2.16, p = 0.04]; Voc12: [t (24) = −2.19, p = 0.038] and Voc8: [t (25) =3.51, p = 0.002]). Task engagement did not alter CEOAE suppression for natural speech [t (24) = 0.62, p = 0.54].

**Figure 2.**
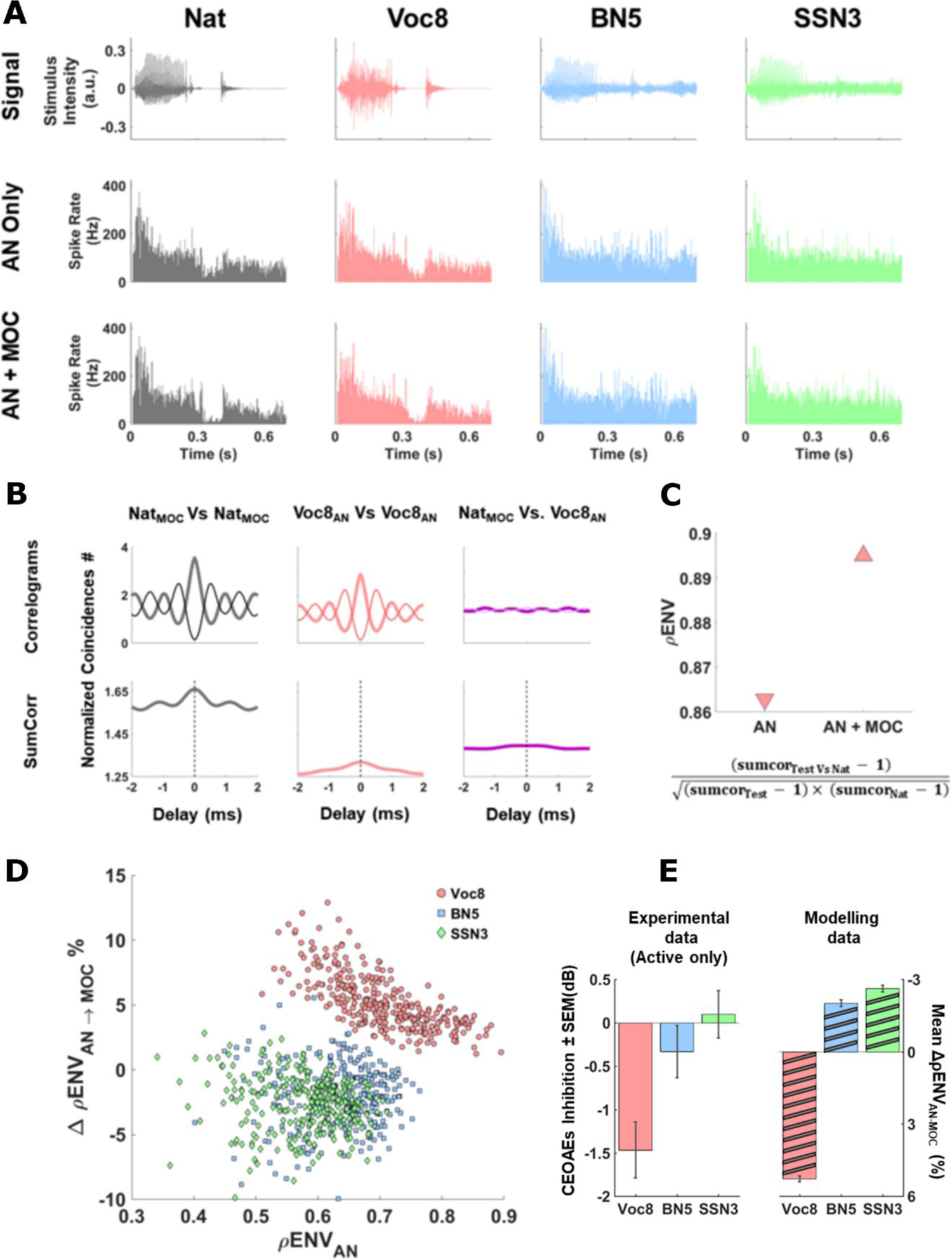
Model auditory periphery output with and without simulated MOC reflex. **A.** Presentation of natural and degraded versions of the word ‘Fuzz’ with and without simulated MOC reflex. ‘Fuzz’ waveforms for natural (*dark gray*, far left), Voc8 (*pink*, second left), BN5 (*light blue*, second right) and SSN3 (*green*, far right) conditions are shown in the top row. Post-stimulus-time-histograms (400 fibers/1ms bin-width) were calculated for low SR AN fibers (Characteristic Frequency: 1kHz) with (bottom row) and without (middle row) simulated MOC reflex. Including simulated MOC reflex reduced activity during quiet for natural condition (and Voc8, but less so) whilst maintaining high spiking rates at peak sound levels (e.g. at 0.075, 0.3 and 0.45ms). No changes in neural representation of signal were visually evident for BN5 and SSN3 ‘Fuzz’. **B & C**. Quantifying ρENV for Voc8 ‘Fuzz’ without simulated MOC reflex. Sumcor plots (bottom row, B) were generated by adding Shuffled Auto-Correlograms (*thick lines*, top left/middle panels, B) and Cross-Correlograms (*thick line*, top right panel, B) to cross-polarity Correlograms (*thin lines*, top row, B) using naturally-spoken ‘Fuzz’ with simulated MOC reflex (left/right columns, B) and Voc8 ‘Fuzz’ without simulated MOC reflex (middle/right columns, B). ρENV for Voc8 ‘Fuzz’ without simulated MOC reflex (AN, *downward-pointing triangle*, C) was calculated from SumCor peaks in B. ρENV with simulated MOC reflex (AN+MOC, *upward-pointing triangle*, C) is also displayed. **D & E**. Comparing ΔρENVs for 300 words after introduction of simulated MOC reflex. Mean percentage changes in ρENVs (calculated across 7 frequencies between 1-2kHz) after adding simulated MOC reflex were plotted against ρENV without simulated MOC reflex for degraded versions of 300 words (each symbol represents one word). ΔρENVs were always positive for Voc8 words (*pink circles*, D) (Max-Min ΔρENV for Voc8: +12.92 to +1.24%), appearing largest for words with lowest ρENVs without simulated MOC reflex. This relationship was absent for BN5 (*light blue squares*, D) and SSN3 (*green diamonds*, D) words whose ΔρENV ranges spanned the baseline (Max-Min ΔρENV for BN5: +5.57 to −9.95%; Max-Min ΔρENV for SSN3: +2.83 to −9.85%). Progression of mean ΔρENVs (± SEM) for model data (*diagonally-striped bars*, right, E) mirrored that of active-task, CEOAE data (mean ± SEM) (*solid-color bars*, left, E).

In contrast, speech in speech-shaped-noise elicited the opposite pattern of results to noise-vocoded speech (Figure 1C). The suppression of CEOAEs was significantly stronger during passive (−0.83 ± 0.26) compared to active listening (−0.16 ± 0.21): rANOVA: [F (1, 24) = 4.44, p = 0.046]. A significant interaction between condition and stimulus-type: [F (2, 48) = 4.67, p = 0.014] confirmed this for both SNRs: (SSN8 [t (27) = 2.71, p = 0.01] and SSN3 [t (28) = 2.67, p = 0.012]) (Figure 1C). Intermediate effects on CEOAEs were observed for words masked by babble noise, with CEOAE significantly smaller than baseline measures during passive, but not active, listening. Cochlear gain was therefore suppressed during active listening of noise-vocoded speech, slightly but significantly suppressed during passive listening in babble noise, and strongly suppressed during passive listening in speech-shaped-noise. Together, our data suggest that the MOC reflex is modulated by task engagement, and strongly depends on the way in which signals are degraded including the type of noise used to mask the speech.

### Auditory brainstem activity reflects changes in cochlear gain when listening to speech-in-noise

The effects of active *vs*. passive listening on the cochlear gain were evident in the activity of subcortical auditory neurons when we simultaneously measured auditory brainstem responses (ABRs) to the same clicks used to evoke CEOAEs.

Click-evoked ABRs—measured during presentation of speech-in-noise—showed similar effects to those observed for CEOAE measurements. Specifically, in both masked conditions wave V— corresponding to neural activity generated in the midbrain by the inferior colliculus (IC) and was significantly enhanced in the active compared to the passive listening condition (Figure 1C) (speech in babble noise: [F (1, 26) = 5.67, p = 0.025] and speech-shaped noise: [F (1, 26) = 9.22, p = 0.005]). No changes in brainstem or midbrain activity were observed between active and passive listening of noise-vocoded speech.

To rule out that the differing results were due to intrinsic differences in the two populations tested (noise-vocoded Vs. masked speech experiments), we compared CEOAE suppression and ABR wave amplitudes in the two groups for active and passive listening of natural speech. No statistical differences were observed for either active or passive listening, therefore, the differences observed in both cochlear gain and auditory brainstem\midbrain activity here can be attributed to the speech degradation presented (active: suppression of CEOAEs [t (23) = −0.21, p=0.83; wave III [t (23) = − 0.45, p=0.65]; wave V [t (23) = 0.09, p=0.93]; passive: suppression of CEOAEs [t (24) = −0.36, p=0.72]; wave III [t (24) = −0.16, p=0.88]; wave V [t (26) = 0.40, p=0.69]).

Together with our CEOAEs results, our data suggests that reduced ABR magnitudes appear only inherited from reduction of the cochlear gain for masked stimuli. These results in the midbrain indicate that while local cochlear gain changes are sufficient when processing single degraded streams such as noise-vocoded speech, processing masked speech requires the involvement of higher-level auditory structures.

### Simulated MOC reflex improves the neural representation of vocoded speech, but not speech-in-noise

To understand why the control of cochlear gain appears to depend on how speech is degraded, we implemented a model of the auditory periphery incorporating MOC reflex through a brainstem circuit with biophysically-realistic temporal dynamics [33]. We specifically tested the hypothesis that suppression of cochlear gain enhances neural encoding of the stimulus envelope via the activation of the MOC reflex. To assess how cochlear gain suppression affects neural representation of speech in AN fibers, natural and degraded (those generating iso-performance in the active task (Voc8; BN5; SSN3)) speech tokens were presented individually to the model at 75 dBA, with and without MOC reflex (Figure 2A). We performed this analysis first for AN fibers with low-spontaneous rate, and high-thresholds given their crucial role in detecting signals in noise as well as their lack of saturation at high intensity levels [34–36].

The neural cross-correlation coefficient, ρENV, a measure of how similar neural envelopes are in different AN spike train responses [37], was employed to quantify the effect of introducing the MOC reflex on envelope encoding at seven logarithmically-spaced frequencies between 1-2 kHz. Values of ρENV, ranging from 0 to 1 for independent to identical neural envelopes respectively, were calculated for the three speech manipulations before and after the MOC reflex was included, with the neural envelope for natural speech acting as the “control” template for comparison. Natural speech “control” simulations were always performed with the MOC reflex, given our observations that a steady CEOAE suppression—indicative of an active MOC reflex— occurred for natural speech experimentally (Figure 1C) and that neural envelopes for natural stimuli were enhanced in model AN fibres with an MOC reflex present (Figure supplement 2).

For the example word, ‘Fuzz’, the three speech degradations in the absence of MOC reflex showed high average similarity in their neural envelopes to their natural speech controls (Figure 2A) (mean ρENV_AN_ for Voc8 = 0.84 ± 0.01; mean ρENV_AN_ for BN5 = 0.65 ± 0.01; mean ρENV_AN_ for SSN3 = 0.63 ± 0.01). However, the addition of the MOC reflex increased the average ρENV significantly for Voc8 (mean ΔρENV = +3.46 ± 0.86%, [t (6) = 10.60, p = 0.0001]); but reduced it significantly for BN5 (mean ΔρENV for BN5 = −3.47 ± 1.88%, [t (6) = −4.88, p = 0.0028]; and also, but not significantly so, for SSN3 (mean ΔρENV for SSN3 −1.42 ± 4.01%, p *=* 0.3859).

Given the range of acoustic waveforms in our speech stimuli, we expanded our analysis to include 300 words (150 stop/non-stop consonants) (Figure 2D). Despite the diversity of speech tokens, the effects of including MOC reflex (on ρENV) were consistently dependent on the type of stimulus manipulation. Neural encoding of speech envelopes improved significantly with simulated MOC reflex for all noise-vocoded words (pink circles, Figure 2D) (mean ΔρENV for Voc8= +5.30 ± 0.12%, [t (299) = 42.71, p = 0.0001]), with the largest enhancements observed for words with the lowest ρENV values in the absence of MOC reflex. In contrast, no such relationship was observed for words in babble noise (BN5, light blue squares, Figure 2D) or speech-shaped noise (SSN3, green diamonds, Figure 2D). Moreover, envelope coding in both speech-in-noise conditions was significantly impaired, on average, when an MOC reflex was introduced (mean ΔρENV for BN5 = − 2.01 ± 0.14, [t (299) = −14.37, p *=* 0.0001]; mean ΔρENV for SSN3 = −2.62 ± 0.12, [t (299) = −22.03, p *=* 0.0001]). This detrimental effect to envelope encoding with MOC reflex was significantly reduced for the masked conditions with higher SNRs (+10 dB SNR for babble noise and +8 dB SNR for speech-shaped noise) (Figure supplement 3B). In contrast, the improvement in neural coding of noise-vocoded speech when introducing MOC reflex was enhanced for stimuli with more noise-vocoded channels (Voc16) (Figure supplement 3B). This suggests that the MOC reflex efferent feedback may specifically be unable to facilitate extraction of signal from noise at specific SNRs.

Given recent evidence that AN fibers with high-spontaneous rate and low-thresholds (fibres that respond preferentially to low intensity sounds but saturate at higher intensities [38–40] may also play an important role in envelope processing [41, 42], we also tested how they processed Voc8, BN5 and SSN3 stimuli with and without MOC reflex. Similarly to AN fibers with high-threshold, we found improvement in the envelope encoding of these low-threshold AN fibres when MOC reflex was included for noise-vocoded speech, but not for the masked conditions (Figure supplement 4), despite the latters’ poorer dynamic range at 75 dBA (normalized sound presentation level across manipulations) likely impacting their overall envelope encoding ability.

The pattern of enhancement of neural envelopes observed across degraded conditions when MOC reflex was implemented in the model (diagonally-striped bars, Figure 2E) mirrored that observed in suppression of CEOAEs for corresponding active listening conditions (solid-colour bars, Figure 2E). Where the MOC reflex was active experimentally for noise-vocoded speech, enhancement of neural envelopes was observed when the same degraded stimuli were presented to the model with MOC reflex present. This was not the case for active listening of masked speech — MOC reflex activity being unaltered experimentally— which was associated with poorer neural representations of the stimulus envelope when MOC reflex was included in the model.

### Cortical evoked potentials reveal enhanced magnitude when active listening to speech-in-noise

The lack of any MOC reflex contribution during active listening to speech masked by speech-like sounds (i.e. babble and speech-shaped noise) compared to noise-vocoded speech suggests other compensatory brain mechanisms must contribute to maintaining iso-performance with noise-vocoded speech. We explored whether higher brain centres—providing top-down perhaps attention-driven, enhancement of speech processing in background noise—contribute to maintaining iso-performance across speech conditions. The significant increase observed in ABR wave V for active speech-in-noise conditions is suggestive of greater activity in the inferior colliculus (IC)—the principal midbrain nucleus which receives efferent feedback from auditory cortical areas. Levels of cortical engagement might then differ across speech manipulations, despite similar overall task performance.

To determine the degree of cortical engagement in the active-listening task, we recorded cortical evoked potentials simultaneously to the CEOAE and ABR measurements from all 56 participants using a 64-channel, EEG-recording system. Grand averages of event-related potentials (ERPs) to speech-onset (Figure 3A and Supplement Figure 5) for the most demanding speech manipulations (Voc8, BN5 and SSN3) were analysed to test the hypothesis that greater cortical engagement occurred when listening to speech-in-noise compared to noise-vocoded speech despite been matched in task difficulty

**Figure 3.**
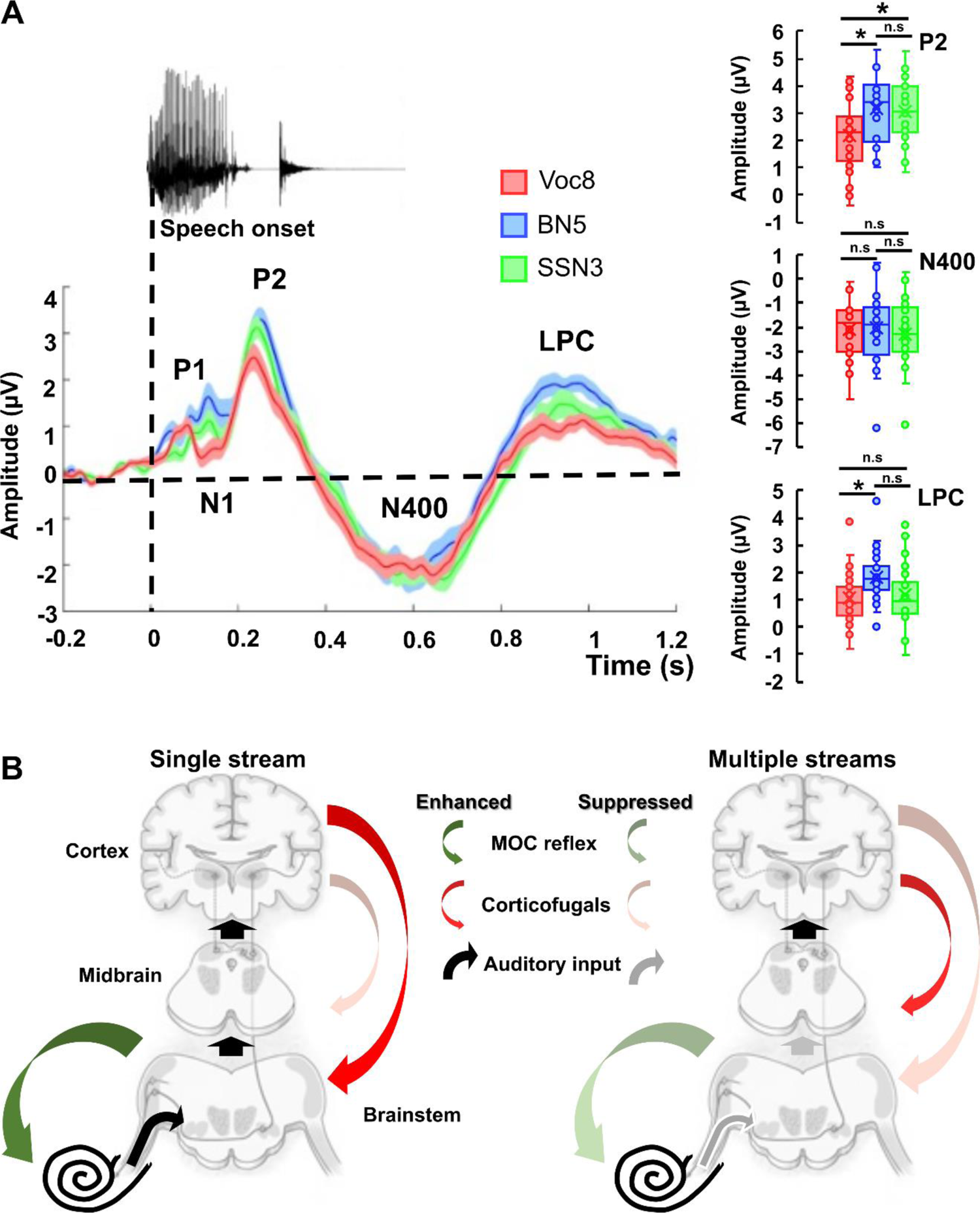
Cortical activity and proposed mechanisms for active listening to noise-vocoded and noise masked speech. **A**. Upper panel shows the “clean” natural spoken word (“Fuzz”) for time-course comparisons with the ERPs. Lower panel shows ERP components (from electrodes: FZ, F3, F4, CZ, C3, C4, TP7, TP8, T7, T8, PZ, P3, P4) during the active listening of Voc8, BN5 and SSN3. Thick lines and shaded areas represent mean and the SEM. Boxplots on the right show statistical comparisons between speech conditions for P2, N400 and LPC components. **B**. Proposed auditory efferent mechanisms for speech processing (Auditory pathway diagrams, [43]. The “Single stream” mechanism shows how isolated degraded tokens such as noise-vocoded speech are processed in a mostly feed-forward manner *(thick black arrows*) (as should be the case for natural speech). The activation of the MOC reflex (*green arrow*) improves the auditory nerve representation of speech-envelope (auditory input) and this information passes up the auditory centres without much need to ‘de-noise’ the signal (represented as black arrows from the cochlea-brainstem-midbrain). Given the trend for increasing cochlear gain suppression with increases in task difficulty, we included the possibility for increased MOC reflex drive from higher auditory regions via corticofugal connections (*red arrow*). In contrast, “multiple streams” such as speech in babble or speech-shaped noise do not recruit the MOC reflex (*shaded green arrow)* because it negatively affects envelope encoding of speech signals (shaded grey arrows from cochlea-brainstem-midbrain). We therefore propose that corticofugal drive is suppressed to the MOC reflex (*shaded red arrow*) leaving greater responsibility for speech signal extraction to the midbrain, cortex and the efferent loop therein (corticofugal connections from auditory cortex to midbrain: *red arrow*). Both mechanisms ultimately lead to equal behavioural performance across speech conditions.

We analysed early auditory cortical responses (P1 and N1 components, Figure 3A) which are largely driven by acoustic features of the stimulus [44, 45]. Noise-vocoded words elicited well-defined P1 and N1 components compared to the speech-in-noise conditions (Figure 3A). This was despite words and noises having similar onsets to noise-vocoded tokens, reflecting the relatively high precision of vocoded speech’s envelope components at stimulus onset compared to those of the masked conditions where the speech and noise envelopes overlap significantly. Later ERP components, such as P2, N400 and the Late Positivity Complex (LPC), have been linked to speech- and task-specific, top-down (context-dependent) processes [44, 46].

Speech masked by babble or speech-shaped noise elicited significantly larger P2 components during active listening compared to the noise-vocoded condition, but not significantly different between themselves [F (2,79) = 5.08, p = 0.008], *post hoc* with bonferroni corrections for 3 multiple comparisons: [BN5 *vs*. Voc8 (p = 0.012); SSN3 *vs.* Voc8 (p = 0.041); BN5 *vs.* SSN3 (p = 1.00)]. Similarly, the magnitude of the LPC—thought to reflect the involvement of cognitive resources including memorization, understanding [47] and post-decision closure [48] during speech processing—differed significantly across conditions: [F (2,79) = 4.24, p = 0.018]. Specifically, LPCs were greater during active listening to speech in babble noise compared to noise-vocoded speech (p = 0.02), with LPCs obtained for active listening to speech in speech-shaped noise intermediate and not significantly different to either (Figure 3A). Consistent with the concept that effortful listening varied across speech-in-noise manipulations even when iso-performance was maintained [49, 50], 2016), the speech manipulation generating the clearest signature of cortical engagement was speech in babble noise, considered the most difficult of the masked conditions (Figure 1B) where iso-performance was achieved at +5 dB SNR (BN5) compared to only +3 dB SNR for speech in speech-shaped-noise (SSN3). In contrast to P2 and the LPC, the N400 component of the ERP— associated with the processing of meaning [51]—did not differ between conditions [F (2,81) = 0.22, p = 0.81]. This is unsurprising, given that participants were equally able to differentiate non-words in Voc8, BN5 and SSN3 conditions, given that iso-performance had been achieved.

Our ERP data are consistent with differential cortical contributions to the processing of noise-vocoded and masked speech, being larger in magnitude for speech manipulations in which the MOC reflex was less efficiently recruited i.e. babble and speech-shaped-noise, and largest overall for the manipulation requiring most listening effort—speech in a background of multi-talker babble. From the auditory periphery to the cortex our data suggests that two different strategies co-exist to achieve similar levels of performance when listening to single undegraded/degraded streams compared to masked speech (Figure 3B). The first involves enhanced sensitivity to energy fluctuations through recruitment of the MOC reflex to generate a central representation of the stimulus sufficient and necessary for speech intelligibility of single streams (Figure 3B, left panel). The second, activated when processing speech in background noise (Figure 3B, right panel), preserves cochlear gain to prevent deterioration of envelope encoding placing the onus of ‘denoising’ on midbrain and cortex including loops between them, to maximise speech understanding.

## Discussion

We assessed the role of attention in modulating the contribution of the cochlear efferent system in a lexical task—detecting non-words in a stream of words and non-words. Employing three speech manipulations to modulate task difficulty—vocoding of words, or words masked by multi-talker babble, or speech-shaped noise (i.e. noise with the same long-term spectrum as speech—we find that these manipulations differentially activate the MOC reflex to modulate cochlear gain. Activation of the cochlear efferent system is also dependent on whether listeners are performing the lexical task (active condition) or are not required to engage whilst watching a silent, stop-motion film (passive condition). Specifically, with increasing task difficulty (i.e. fewer noise-vocoded channels), noise-vocoding increasingly activates the MOC reflex in active, compared to passive, listening. The opposite is true for the two masked conditions, where words presented at increasingly lower SNRs more strongly activate the MOC reflex during passive, compared to active, listening. By adjusting parameters of the three speech manipulations—the number of noise-vocoded channels, or the SNR for the speech-in-noise conditions, we find that lesser MOC reflex activity is accompanied by greater magnitude of cortical activation, to maintain iso-performance in the task. A computational model incorporating efferent feedback to the inner ear demonstrates that improvements in neural representation of the amplitude envelope provides a rationale for either suppressing or maintaining the cochlear gain during the perception of noise-vocoded speech or speech-in-noise respectively. Our data suggest that a network of brainstem and higher brain circuits is involved in maintaining performance in an active listening task, and that different aspects of this network, including reflexive circuits in the lower brainstem, and the relative allocation of attentional resources, are differentially invoked depending on specific features of the listening environment.

### Attentional demands reveal differential recruitment of MOC reflex

Our data highlight a categorical distinction between active and passive processing of single, degraded auditory streams (vocoded speech) and parsing a complex acoustic scene to hear out a stream from multiple competing, spectrally-similar sounds (multi-talker babble and speech-shaped noise). Specifically, task difficulty during active listening appears to modulate the cochlear gain in a stimulus-specific manner. The reduction in cochlear gain with increasing task difficulty for noise-vocoded speech and, conversely, the preservation of cochlear gain when listening to speech in background noise, suggests that attentional resources might gate the MOC reflex differently depending on how speech is degraded. In contrast to active listening, where participants were asked to ignore the auditory stimuli and direct attention to a silent film, the MOC reflex was gated in a direction consistent with the auditory system suppressing irrelevant and expected auditory information whilst (presumably) attending to visual streams [52–54]. Interestingly, activation of the MOC reflex was observed for natural speech—further evidence that activation is not limited to tones and broadband noise [55–57]—and did not depend on whether participants were required to attention (i.e: were engaged in a lexical-decision task) or not. This can be explained by natural speech being particularly salient as an undegraded, ethologically-relevant stimulus and the low attentional load of passively watching a film resulting in the continued monitoring of unattended speech [58].

To explain the variable reduction in cochlear gain between noise-vocoded and masked stimuli across active and passive conditions, top-down and bottom-up mechanisms can be posited as candidates to modify the activity of the MOC reflex. Bottom-up mechanisms, such as the increased activation of the MOC reflex by wideband stimuli with flat power-spectra [59] may explain a facet of our results—especially during passive listening. For instance, reduced activation of the reflex observed for babble noise during passive listening may arise from being a weaker suppressor than ‘stationary’ noises such as white noise, or speech-shaped noise [60, 61]. However, noise-vocoded stimuli with relatively fewer channels might have also been thought to have activated the MOC reflex more effectively due to their more ‘noise-like’ spectra. The absence of suppression of CEOAEs in the passive noise-vocoded conditions, as well as the pattern of activation of the MOC reflex across all active listening conditions, suggests that a perceptual, top-down categorization of the stimuli is necessary to appropriately engage or disengage the MOC reflex.

### Descending control of the MOC reflex for speech stimuli is likely bilateral

A central premise of our study, and those exploring the effects of attention on the MOC reflex, is that otoacoustic emissions recorded in one ear can provide a direct measure of top-down modulation of cochlear gain in the opposite ear. However, it has also been suggested that activation of the MOC reflex may differ between ears to expand interaural cues associated with sound localization [62]. This process could be independently modulated by the extensively ipsilateral corticofugal pathways described anatomically (see [63] for review). Had the activation of the MOC reflex been independently controlled at either ear—for example to suppress irrelevant clicks in one ear whilst preserving cochlear gain in the ear stimulated by speech—we would have expected similar suppression of CEOAEs across active and passive conditions for all speech manipulations—given that click stimuli were always irrelevant to the task. Instead, however, the suppression of CEOAEs was both stimulus- and task-dependent, reducing the likelihood that dichotic stimulus presentation engaged top-down modulation of cochlear gain differentially at either ear. It remains unclear whether spatial hearing, and any role the MOC reflex plays therein, impacts top-modulation given recent evidence of weaker spatial segregation cortically in tasks requiring high perceptual demand (as was the case at iso-performance for both noise-vocoded and masked speech conditions in our study) [64].

Anatomical evidence of purely ipsilateral corticofugal pathways ignores the possibility that, even when presented monaurally, descending control of the MOC reflex for speech stimuli may likely be bilateral. Unlike pure tones, speech activates both left and right auditory cortices even when presented unilaterally to either ear [65]. In addition, cortical gating of the MOC reflex in humans does not appear restricted to direct, ipsilateral descending processes that impact cortical gain control in the opposite ear [66]. Rather cortical gating of the MOC reflex likely incorporates polysynaptic, decussating processes that influence/modulate cochlear gain in both ears.

### Stimulus-specific enhancement of speech envelope’s neural coding explains observed variability in the suppression of CEOAEs

Beyond extending spatial hearing [62] and the protection from noise exposure [67–69], the function most commonly attributed to the activation of the MOC reflex has been its ‘unmasking’ of transient acoustic signals in the presence of background noise [17,55,70]. By increasing the dynamic range of mechanical [71, 72] and neural [16, 17] responses to amplitude-modulated components in the cochleae [15, 73], it has been suggested that suppression of cochlear gain by the MOC reflex might preferentially favour the neural representation of syllables and phonemes in speech over masking noises.

Whilst models of the auditory periphery that include efferent feedback, typically demonstrate improved word recognition for a range of masking noises (pink noise [74–76], speech-shaped noise [77] and babble noise [75]), positive correlations between increased activation of the MOC reflex and improved speech-in-noise perception are only sporadically reported [23, 27], with some studies reporting negative [22, 28] or non-significant correlations [29, 30]. This variability has generally been attributed to methodological differences in OAE measurement (see [78] for review). Still, more recent studies have questioned whether suppression of cochlear gain by the MOC reflex actually benefits hearing in noise: proposing either a contribution of the reflex to perception only at specific SNRs [24, 78] or, *in extremis*, that the MOC reflex plays no role, and that neural adaptation to noise statistics may instead underlay robust recognition of degraded speech [73,79,80].

However, the stimulus- and task-dependent suppression of cochlear gain we report when speech was simultaneously employed as a contralateral activator of the MOC reflex and as a target in a lexical-decision task—rather than performing sequential experiments to correlate the strength of the reflex as function of speech-in-noise performance, as in the majority of the reports— suggests that the automatic activation of the MOC reflex is gated on or off by top-down modulation depending on whether it functionally benefits or not performance. Our modelling data support this conclusion by accounting for the stimulus-dependent suppression of click-evoked OAEs through enhancement of speech-envelope encoding in the auditory nerve. The apparent benefit of suppressing cochlear gain to neural coding of the envelope of noise-vocoded words, compared to a disbenefit for words masked by background noise, is consistent with noise-vocoded words retaining relatively strong envelope modulations, and that these modulations are extracted effectively through expansion of the dynamic range as cochlear gain is reduced. For both noise-maskers, however, any improvement to envelope coding due to dynamic range expansion applies to both speech and masker, since these overlap spectrotemporally, resulting in poorer representation of the speech envelope. Whilst the results of our simulations are based on average changes in the neural correlation coefficient with efferent feedback calculated for 300 words, individually, the effect of suppressing cochlear gain was highly dependent on both the choice of word and the SNR at which it was presented (Figure 2C and Figure supplement 3). If top-down modulation does indeed gate the activation of the MOC reflex, it must account for the statistical likelihood that suppression of cochlear gain can improve the neural coding of the stimulus’ envelope over the duration of the task. One potential criterion for predicting whether activation of the MOC reflex can improve speech intelligibility is the extent to which target speech is ‘glimpsed’ in spectrotemporal regions least affected by a background noise [81–83]. If computing speech envelope SNR in short time windows is indeed key to speech intelligibility—as proposed by several models [84, 85]— then this could be a suitable metric by which top-down modulation of the MOC reflex could be adjusted, explaining why previous studies of OAE suppression, that employ identical paradigms aside from the stimuli with different spectrotemporal content (e.g: consonant-vowel pairs), can generate inconsistent correlations between discrimination-in-noise and the strength of the MOC reflex [22, 86]. Additionally, changes in speech envelope SNR as a result of presenting words at different SNRs should impact any benefit of activating the MOC reflex to speech intelligibility, offering an explanation for correlations previously reported between the strength of the MOC reflex and task performance at varying SNRs [24, 78].

In our model, switching from lowest to intermediate stimulus SNRs for both noise-maskers led to an increase in the average ΔρENV when efferent feedback was applied (Supplemental Figure 3B). However, values of ΔρENV at the intermediate SNRs were positive for only a minority of words tested (73 SSN8 words and 116 BN10 words, Supplemental Figure 3A). This is consistent with the lack of OAE suppression observed experimentally in the active tasks for babble noise at 10 dB SNR and speech shaped noise at 8 dB SNR (Figure 1C). Where an active MOC reflex does not functionally benefit the neural representation of the speech envelope, it remains possible that it is activated in another capacity, for example to prevent damage by prolonged exposure to loud sounds [67–69].

### Limitations of current auditory periphery model and sumcorr analysis

As a stimulus feature strongly correlated with the cortical tracking and decoding of speech [87–89] and the basis of several successful speech intelligibility models [84, 85], the speech envelope appears central to our understanding of speech. Nevertheless, the relative contributions of envelope and temporal fine structure (TFS) cues to speech intelligibility remain to be determined [85,90–94]. Consequently, future investigations of the functional role of the MOC reflex may benefit from valid comparisons of the neural envelope and TFS representations of stimulus in response to degraded speech. Such comparisons have been performed previously for noise-vocoded stimuli employing a simplified model of the auditory periphery with lateral inhibition [92].

Quantifying sumcorr peaks served well here as a metric to correlate neural envelopes of natural and differentially degraded speech. While this analysis highlighted the similarity between natural and noise-vocoded speech envelopes, its results for our noise-masker simulations appear counterintuitive, especially given that studies applying the same model as here, but with basic efferent feedback, have previously suggested a role for the MOC reflex in speech-in-noise discrimination [74–76]. These studies have employed an automatic speech recognizer (ASR) to analyse AN output, encoding the shape of the speech spectrum in brief time windows to estimate the speech token most likely presented as the stimulus. However, their performance reduces for natural speech as efferent feedback increases [74], suggesting that this analysis would not account for the experimentally observed activation of the MOC reflex for natural speech. Nevertheless, further assessment of the two analyses’ relative validity remains difficult, given the number of model parameters that differed between this and prior studies (frequency range sampled, speech tokens presented, their stimulus levels and how efferent feedback was implemented).

As regards the auditory periphery model itself, recent insights into the MOC reflex’s circuitry will enhance the biological realism of any future models incorporating efferent feedback. Our model and previous phenomenological models simulate efferent feedback as a solely on-frequency effect which appears unlikely given anatomical evidence that MOC neurons are sparse and innervate relatively broad regions of the cochlea [95, 96]. Additionally, most models do not include neural adaptation to stimulus statistics as a potential contributing factor to speech-in-noise discrimination [73, 79], we therefore cannot discount its involvement in the lexical-decision task.

### Higher auditory centre activity supports co-existence of multiple strategies to achieve similar levels of performance

The impact of modulating the MOC reflex was observed in the activity of the auditory midbrain and cortex. Increased midbrain activity for active noise-masked conditions was consistent with changes in magnitudes of ABRs previously reported during unattended Vs attended listening to speech [97] or clicks [98, 99]. This highlights the potential for subcortical levels to either enhance attended signals, or filter out distracting auditory information. At the cortical level, recorded potentials were larger for all attended compared to ignored speech (Supplemental Figure 5), consistent with previous reports [100, 101]. However, later cortical components were of greater magnitude for masked than noise-vocoded speech while attending to the most extreme manipulations. Late cortical components have been associated with the evaluation and classification of complex stimuli [46] as well as the degree of mental allocation during a task [44]. Therefore, differing cortical activity likely reflects greater reliance on, or at least increased contribution from, context-dependent processes for speech masked by noise than for noise-vocoded speech.

Taken together, the observed differences in physiological measurements from higher auditory centres and the auditory periphery highlight the possibility of diverging pathways to process noise-vocoded and masked speech. Evidence for systemic differences in processing single degraded streams, compared to masked speech, has been reported in the autonomic nervous system [49]: where, despite maintaining similar task difficulty across conditions, masked speech elicits stronger physiological reactions than single unmasked streams. Here we propose that two strategies allow maintaining iso-performance even when stimuli are categorically different. The processing of single and intrinsically degraded streams selectively recruits auditory efferent pathways from the auditory cortex to the inner ear which in turns improves the representation —‘de-noise’—of the stimuli in the periphery (Figure 3B). In contrast, multiple streams appear to rely much more on higher auditory centres such as midbrain and auditory cortex for the extraction of foreground, relevant signals (Figure 3B). Evidence of de-noising auditory signals in the cortex [87,102–104] has led to the oversight of the potential role of the extensive loops and chains of information between cortex, subcortical regions and the auditory periphery in everyday listening environments [20,105–107] and therefore as target for hearing technologies.

### Implications for hearing-impaired listeners

Although normal-hearing listeners appear to benefit from an MOC reflex that modulates cochlear gain, and is amenable to top-down, attentional control, it is important to note that users of cochlear implants (CIs)—for whom normal-hearing listeners processing noise vocoded speech have often been a proxy—have no access to the MOC reflex. CIs bypass the mechanical processes of the inner ear, including the outer hair cells—which are non-functioning or absent completely in individuals with severe-to-profound hearing loss—to stimulate directly the auditory-nerve fibers themselves. Efforts have been made to incorporate MOC-like properties into CI processes, at least to provide for expanded auditory spatial cues to improve listening in bilaterally implanted CI users [62], but the capacity to exploit efferent feedback to aid speech understanding in CI listeners is yet to be exploited in any device.

For other hearing-impaired listeners, aided or unaided, the contribution of MOC feedback to speech processing is limited compared to normal-hearing listeners as, in most cases, their hearing loss comes from damage to the OHCs, which receive direct synaptic input from MOC fibers. In hearing loss generally, the degradation or loss of peripheral mechanisms contributing to effective speech processing in complex listening environments may mean that listeners rely more heavily on attentional and other cortical-mediated processes, contributing to widely reported increases in listening effort required to achieve adequate levels of listening performance [50]. This increase in listening effort—likely manifesting over time—may not be reflected in performance in relatively short, laboratory- or clinic-based assessments of hearing function.

## Materials and Methods

This study was approved by the Human Research Ethics Committee of Macquarie University (ref: 5201500235). Each participant signed a written informed consent form and was provided with a small financial remuneration for their time.

### Hearing assessment

All 66 participants (36 females, aged between 18-35 (mean: 24 ± 7 years old)) included in this study had normal pure tone thresholds (< 20 dB HL); normal middle ear function (standard 226 Hz tympanometry; normal OHCs function (assessed with Distortion Product OAEs (DPOAEs) between 0.5-10 kHz. Steady-state contralateral and ipsilateral broadband noise middle ear muscle reflexes (MEMR) were assessed and all participants had thresholds > 75 dB HL.

### Experimental Protocol

Participants were seated comfortably inside a sound-proof booth (ISO 8253-1:2010) while wearing an EEG cap (Neuroscan 64 channels, SynAmps2 amplifier, Compumedics Limited). Two listening conditions (passive and active) were counterbalanced across participants. In the passive listening condition, subjects were asked to ignore the auditory stimuli and to watch a non-subtitled, stop-motion movie. To ensure participants’ attention during this condition, they were monitored with a video camera and were asked questions at the end of this session (e.g. What happened in the movie? How many characters were present?). The aim of a passive or an auditory-ignoring condition is to shift attentional resources away from the auditory scene and towards the visual scene. During active listening, participants performed an auditory lexical-decision task, where they were asked to press a keyboard’s space key each time they heard a non-word in strings of 300 speech tokens. d’ was used as a measure of accuracy and calculated as: Z(correct responses) – Z(false alarm).

Simultaneous to the presentation of word/non-word in one ear, CEOAEs were recorded continuously in the contralateral ear (Figure 1A). The ear receiving either the clicks or speech stimuli was randomized across participants.

### Speech Stimuli

423-word items were acquired from Australian-English-adapted versions of monosyllabic consonant-nucleus-consonant (CNC) word lists [108] and were spoken by a female, native Australian-English speaker. The duration of words ranged between 420-650 ms. 328 monosyllabic CNC non-word tokens were selected from the Australian Research Council non-word database [109]. Speech stimuli were delivered using ER-3C insert earphones (Etymotic Research, Elk Grove Village, IL) and Presentation® software (Neurobehavioral Systems, Inc., Berkeley, CA, version 18.1.03.31.15) at 44.1 kHz, 16-bits. Blocks of 200 words and 100 non-words (randomly selected from the speech corpus) were presented for each of the speech manipulations (see details below). Speech tokens were counterbalanced in each block based on the presence of stop and non-stop initial consonants: 100 stop/ non-stop consonant words; 50 stop/non-stop consonants words with a maximum of 3 repeats per participant allowed. Each block had a duration of 12 minutes, (Figure 1A) and participants could take short breaks between them if needed. The order of blocks was randomized to prevent presentation order bias or training effects [110].

### Noise-vocoded Speech

Twenty-seven native speakers of Australian-English (17 females: 25 right-handed, 2 left-handed) were recruited (aged between 18-35 (mean: 23 ± 5 years old). Based on the noise vocoding method and behavioural results of Shannon et al. [111], three noise-vocoded conditions (16-, 12- and 8-channels: Voc16, Voc12, and Voc8, respectively) were tested to represent 3 degrees of speech intelligibility (i.e. task difficulty). Four blocks were assessed in both active and passive listening conditions: Block 1: natural speech; Block 2: Voc16; Block 3: Voc12 and Block 4: Voc8, for a total duration of 2.6 hours (including hearing assessment and EEG cap set-up).

### Speech in Babble Noise

Twenty-nine native speakers of Australian-English (19 females: 28 right-handed, 1 left-handed) were recruited, aged between 20-35 (mean: 26 ± 9 years old). The BN used consisted of four females and four male talkers [112] and was filtered to match the long-term average spectrum of the speech corpus. Random segments from 60 secs BN recording were selected as noise maskers. Both speech tokens and the BN segments were matched in duration, Figure supplement 6. Three blocks were presented in the active and passive listening conditions: Block1: natural speech; Block 2: BN at +10 dB SNR (BN10) and Block 3: BN at +5 dB SNR (BN5).

### Speech in Speech-Shaped noise

SSN was generated to match the long-term average spectrum long-term average of the speech corpus, Figure supplement 6. Segments from 60 secs SSN were randomly selected as noise maskers. Both BN and SSN manipulations were presented in the same session, therefore, Block1 was the same for both manipulations, Block 2: SSN at +8 dB SNR (SSN8) and Block 3: SSN at +3 dB SNR (SSN3). A total of 10 blocks (5 blocks in the active and 5 blocks in the passive condition) of 12 min duration were presented to the 29 subjects who participated in BN and SSN experiments. All blocks were presented in a unique session that lasted 3 hours. All tokens were root-mean-square normalized and the calibration system (sound level meter (B&K G4) and microphone IEC 60711 Ear Simulator RA 0045 563 (BS EN 60645-3:2007), (see CEOAEs acquisition and analysis section)) was set to 75 dB “A-Weighting” which matches the human auditory range.

### CEOAEs acquisition and analysis

Non-filtered click stimuli, with a positive polarity and 83 µsec of duration were digitally generated using RecordAppX (Advanced Medical Diagnostic Systems) software. The presentation rate was 32 Hz in all conditions which contributed to minimize ipsilateral MOCR activation [113]. Ipsilateral MOCR activation was otherwise constant across participants and experimental manipulations by maintaining a fixed click-rate.

Both the generation of clicks and OAE recordings were controlled via an RME UCX soundcard (RME, Haimhausen, Germany), and delivered/collected to and from the ear canal through an Etymotic ER-10B probe connected to ER-2 insert earphones with the microphone pre-amplifier gain set at 20 dB. Calibration of clicks was performed using a sound level meter (B&K G4) and microphone IEC 60711 Ear Simulator RA 0045 (BS EN 60645-3:2007). This setup was also used to calibrate the speech stimuli. In addition, clicks were in-ear calibrated using forward equivalent pressure level (FPL) ensuring accurate stimulus levels [114]. The OAE’s probe was re-positioned, re-calibrated and the block re-started if participants moved or touched it.

CEOAE data were analysed off-line using custom Matlab scripts (available upon request) [113]. The averaged RMS magnitudes of CEOAE signals were analysed between 1-2 kHz given maximal MOC effects in this frequency band [115]. Only binned data for averaged CEOAEs displaying a SNR ≥ 6 dB and with > 80% of epochs retained (i.e. had RMS levels within the two standard deviations limit), were selected as valid signals for further analysis. The initial and final 60 secs of CEOAE recordings of each block, in the absence of speech tokens (Figure 1A), were used as baseline. As no significant differences were observed between CEOAE baseline magnitudes within participants (p > 0.05) across experimental conditions, all baselines were pooled within participants. This allowed for an increase in both SNR and reliability of the CEOAEs individually. After baseline recording, CEOAEs were continuously obtained for 10 minutes during the presentation of the speech tokens. The inhibition of CEOAE magnitude relative to the baseline was calculated as follows and reported as means and standard errors of the mean: CEOAE inhibition = CEOAE_speech presentation (average across minutes)_ – CEOAE_baseline (first 60 sec)_

### Stimuli calibration

Controlling the stimulus level is a critical step when recording any type of OAE due to the potential activation of the middle ear muscle reflex (MEMR). High intensity sounds delivered to an ear can evoke contractions of both the stapedius and the tensor tympani muscles causing the ossicular chain to stiffen and the impedance of middle ear sound transmission to increase [116, 117]. As a result, retrograde middle ear transmission of OAE magnitude can be reduced due to MEMR and not MOCR activation [118]. For this reason, we were particularly careful to determine the presentation level of our stimuli. The presentation level for all natural, noise-vocoded and speech-in-noise tokens was 75 dBA and click stimulus at 75 dB p-p, no MEMR contribution was expected given a minimum of 10 dB difference between MEMR thresholds (see Hearing Assessment section) and stimulus levels (ANSI S3.6-1996 standards for the conversion of dB SPL to dB HL. All tokens were root-mean-square normalized calibrated using the same CEOAEs calibration system (see **CEOAEs acquisition and analysis**).

### EEG acquisition and analysis

EEG measurements and the CEOAE setup were synchronized using a Stimtracker (https://cedrus.com/). EEG data were acquired according to the 10–20 system. Impedance levels were kept below 5 kΩ for all electrodes. Signals were sampled at a rate of 20 kHz in the AC mode with a gain of 2010 and an accuracy of 0.15 nV/LSB. Early and late ERP components were analysed offline using Fieldtrip-based scripts [119]. Data were re-referenced to the average of mastoid electrodes. Trials started 200 ms before and ended 1.2 sec after speech onset. Components visually identified as eye blinks and horizontal eye movement were excluded from the data as well as trials with amplitude >75 µV. The accepted trials (60-80% per condition) were band-pass filtered between 0.5 to 30 Hz with transition band roll-off of 12 dB/octave. Trials were baseline-corrected using the mean amplitude between −200-0 ms from speech onset. Baseline-corrected trials were averaged to obtain ERP waveforms. Analysis windows centred on the grand average-ERP component maximums were selected [120]: P1 (100-110 msec) and N1(145-155 msec); P2 (235-265 msec), N400 (575-605 msec) and LPP (945-975 msec). Mean amplitude for each component within the analysis window was calculated for each participant and experimental condition.

### EEG. Auditory Brainstem Responses (ABRs)

ABR signals were extracted from central electrodes (FZ, FCZ, CZ). 15 msec-duration ABR analysis windows (5 ms prior and 10 ms after click onset) were selected. A total of 19200 trials (click rate of 32 Hz across 10 minutes per block) were band-pass filtered between 200-3000 Hz. Averaged ABR waveforms were obtained using a weighted-averaging method [121]. Amplitude of waves III and V (Figure 1A) were visually determined for each subject across blocks and conditions when appropriate (wave amplitudes above the residual noise, therefore a positive SNR). Due to stimulus level restrictions (<= 75 p-p dB SPL to avoid MEMR activation), wave I could not be extracted from the EEG residual noise.

### Statistical Analysis

Sample size estimation was computed according to the statistical test employed by using G*Power [122] (Effect size f = 0.4; α err prob = 0.05; Power (1-β err prob) = 0.8). All variables were tested for normality (Shapiro-Wilk test), outlier residual values preventing normal distribution were removed from the data set (Table supplement 2 and 3). One-way ANOVA for the behavioural and ERPs data; repeated measures ANOVAs (rANOVA) for CEOAEs and ABRs data and t-tests (alpha=0.05, with bonferroni corrections for multiple comparisons) were performed. One-way ANOVAs had Stimuli-Type (Natural, Voc16, BN10…etc) as Factor whereas rANOVA had both Conditions (Active and Passive) and Stimuli-type as Factors. The interaction (Conditions x Stimuli) was also explored.

### Auditory Nerve and Brainstem Simulations

The Matlab Auditory Periphery and Brainstem (MAP-BS) computational model was used to simulate auditory nerve (AN) responses with and without an efferent feedback loop at seven frequencies (1000 Hz, 1122 Hz, 1260 Hz, 1414 Hz, 1587 Hz, 1782 Hz, 2000 Hz). A binaural brainstem circuit (Figure supplement 1A) using Hodgkin-Huxley models was implemented to calculate within-channel efferent attenuation of the cochlear gain with biophysically realistic temporal dynamics [33] (Figure supplement 1B). The current balance equation for all single-compartment, Hodgkin-Huxley models was described as follows:

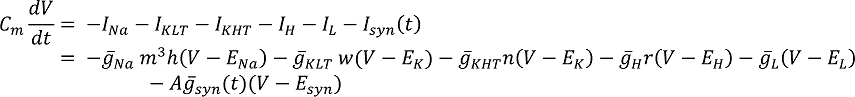

Where *^g̅^ x* is the maximum channel conductance and *Ex* is the corresponding reversal potential for conductance *x* where *x* was either: *Na* (fast sodium), *KLT* (low threshold potassium), *KHT* (high threshold potassium), *H* (hyperpolarization-activated mixed-cation), *L* (leak) and *K* (all potassium conductances) (see Table supplement 1 for relative values in cell types). *^g̅^ syn* (*t*) is the unitary synaptic input at time, *t*, and *A* is an input activity-dependent factor with a predetermined, maximum value for each cell type (*Max Synaptic Current*, Table supplement 1). *Esyn* represents the synaptic reversal potential which was either 0mV for excitation or −70mV for inhibition (Figure supplement 1A and Table supplement 1). An absolute voltage threshold of −20mV was used for suprathreshold responses. The dynamics of gating variables (*m*, *h*, *w*, *n* and *r*) were described by the following differential equation:

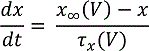

Where *τx(V)* is the time constant of the variable *x* and *x∞(V)* is the steady state value of *x* at voltage, *V*. See ‘HH5lookupInt.m’ file in MAP-BS [33] for values of *τx(V)* and *x∞(V)* for individual conductances. The spiking output of MOC neurons was used to calculate inhibition of the basilar membrane displacement’s nonlinear path [75] as follows:

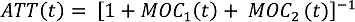

Where the formula for calculating activation and exponential decay of *MOCx* at time, t, was expressed as:

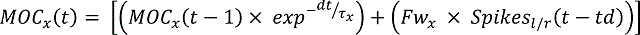

Where τx is the decay time constant (55ms for *MOC1* and 400 ms for *MOC2* (Backus and Guinan, 2006) and *Spikes(t-delay)* is spikes observed across ‘left’ and ‘right’ MOC populations at a time, *t-td* (where *td* equals a 10ms lag associated with synaptic latencies from MOC fibers to OHCs [123]. *F* is a temperature- and sample rate-dependent factor attenuating *Spikes(t-delay)* appropriately (0.02 here for a default temperature of 38 C and a sample rate of 100 kHz). *wx* is a weighting factor determining the relative contribution of MOC1 and MOC2 to attenuation (0.9 and 0.1 for MOC1 and MOC2 respectively).

### Word Presentation

300 words (150 stop/non-stop consonant words), were chosen at random from the speech corpus and were degraded using the most demanding speech manipulations (Voc8, BN5 and SSN3). Normal, Test+ and polarity-inverted, Test-, versions of each manipulation were presented to the MAP-BS model at 75 dB SPL both with and without efferent feedback. Natural words (both normal, Nat+, and polarity-inverted, Nat-) were also presented to the MAP-BS model with efferent feedback enabled as a reference AN output (Figure supplement 2).

### Shuffled Auto-Correlogram Analysis

Comparative analysis of AN coding of amplitude-modulated (AM) envelope between Voc8/BN5/SSN3 conditions and the reference natural condition was performed using shuffled auto- and cross-correlograms (SACs and SCCs respectively)[124] Normalized all-order histograms were calculated using the spike trains of 400 low SR, AN fibers with a coincidence window of 50µs and a delay window ± 25 ms centred on zero [37]. No correction for triangular shape was required given brevity of delay window relative to stimulus length (between 420-650 ms) [124]. A neural cross-correlation coefficient, *ρENV*, quantifying AM envelope encoding similarity between conditions was generated as follows [37]:

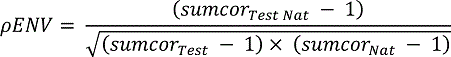

Where sumcor_Nat_ (natural word reference) and sumcor_Test_ (Voc/BN/SSN conditions) are the averages of SACs (Normalized all-order histograms for Nat+ *vs.* Nat+/Test+ *vs.* Test+ for sumcor_Nat_/sumcor_Test_ respectively) and cross-polarity histograms (Normalized all-order histograms for Nat+ *vs.* Nat-/Test+*vs.*Test- for sumcor_Nat_ /sumcor_Test_ respectively). Sumcor_Test Nat_ is the average of the SCC (Average of normalized all-order histograms for Nat+ *vs.* Test+ and Nat-*vs.* Test-) and the cross-polarity correlogram (Average of normalized all-order histograms for Nat-*vs.* Test+ and Nat+ *vs*. Test-) between natural and Voc8/BN5/SSN3 conditions. All high-frequency oscillations (> test-frequency), associated with fine-structure leakage, were removed from sumcors (Heinz and Swaminathan, 2009). *ρENV* values ranged from 0 to 1 where 0 represents completely dissimilar spike trains and 1 represents identical spike patterns [37].

### Analysis of modelling and statistics

Percentage changes in *ρENV* due to efferent feedback inclusion in MAP-BS were calculated for each test frequency and Voc8/BN5/SSN3 condition as follows:

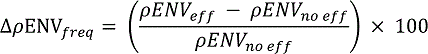

Where Δ*ρENV_freq_* is the percentage change in *ρENV* at a test-frequency for a manipulated word. *ρENV_no eff_* and *ρENV_eff_* are measures of *ρENV* with and without efferent feedback enabled respectively. An average Δ*ρENV_freq_* was calculated across test-frequencies for each word and manipulation. A one-sample t-test was then performed to confirm whether average Δ*ρENV_freq_* for all words differed from zero for each speech manipulation. Data are reported as means and standard errors of the mean.

## Acknowledgements

H.H.P. was supported in this study by an International Macquarie University Excellence Scholarship and the Hearing CRC. The authors thank Prof. David Poeppel for his contributions during experimental design. We thank Prof. David Ryugo for his comments on the manuscript. In addition, we thank Ronny Ibrahim, Jaime Undurraga and Greg Stewart for their technical support regarding equipment and EEG analysis; Andrew Brughera for his help with figures and Nicholas Clark for his assistance with MAP-BS.

## Competing interests

The author has no financial interests to declare. The other authors declare that no competing interests exist.

## Supplemental Information

**Figure supplement 1.**
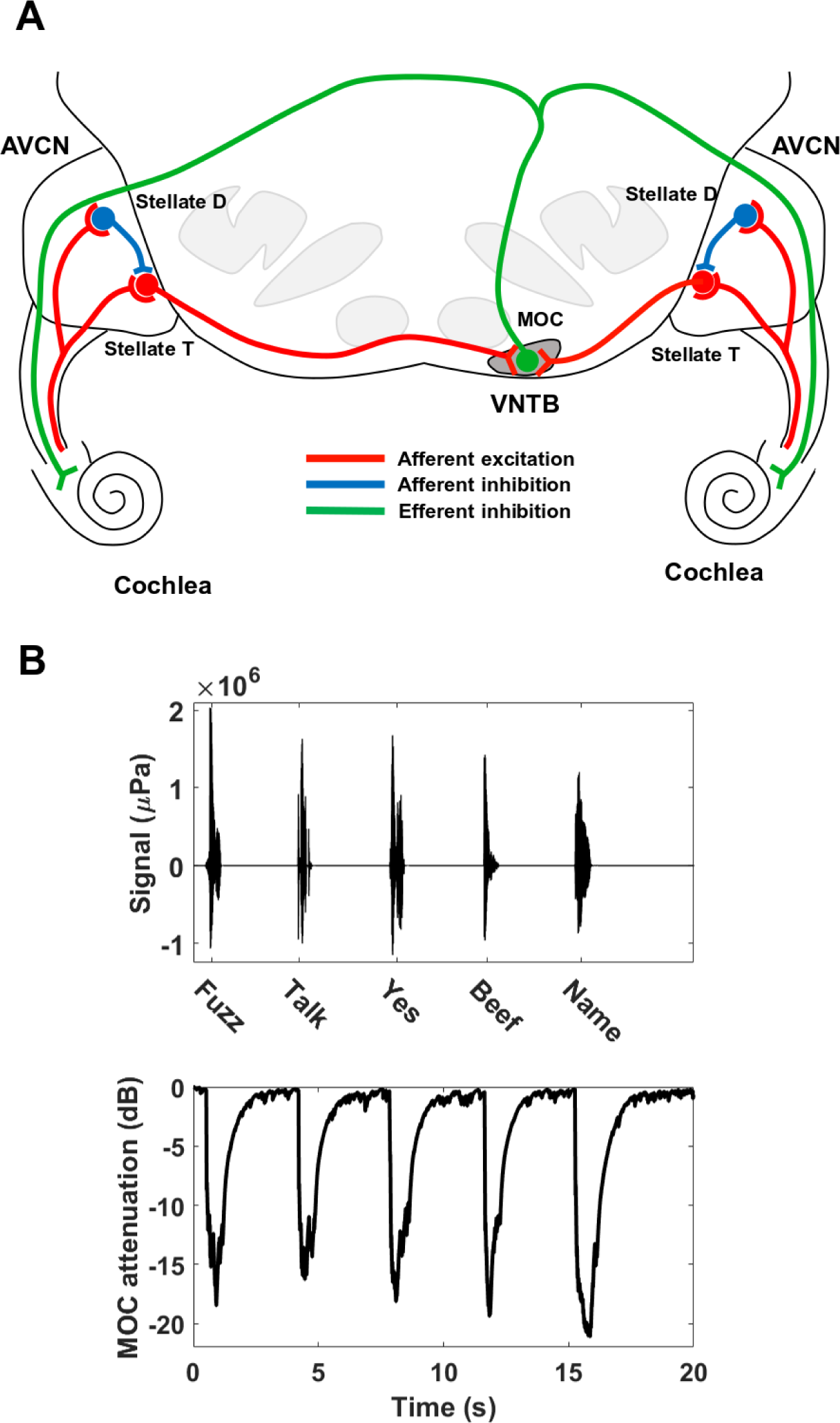
Brainstem circuit and MOC attenuation of MAP-BS. **A**. Overview of brainstem circuit in MAP-BS. Bilateral afferent excitation (*red lines*) via high spontaneous rate auditory nerve (AN) fibers drives Stellate T (within-channel AN input) and D neurons (wideband AN input) in anteroventral cochlear nucleus (AVCN). Stellate D neurons provide ipsilateral, feedforward inhibition to Stellate T neurons in the same frequency channel (*blue lines*). Ventral nucleus of the trapezoid body (VNTB) neurons receive bilateral excitation from Stellate T neurons; their output is combined across ‘left’ and ‘right’ brainstems to calculate within-channel, efferent attenuation to both cochleae (*green lines*). Emulating inhibition of cochlear amplification, VNTB output impacts the non-linear pathway of dual-resonance nonlinear model of cochleae with a short delay [75, 123]. **B**. Temporal dynamics of efferent feedback in MAP-BS. Upper panel shows waveforms of naturally-spoken word tokens presented diotically at 75 dB to MAP-BS model with a 3 sec pause between tokens as in the psychoacoustic paradigm (Figure 1A). Lower panel presents MOC attenuation calculated across ‘left’ and ‘right’ VNTB populations. MOC attenuation did not summate across series of word tokens but recovered to baseline; single-shot presentation of word tokens was therefore implemented for the main results.

**Figure supplement 2.**
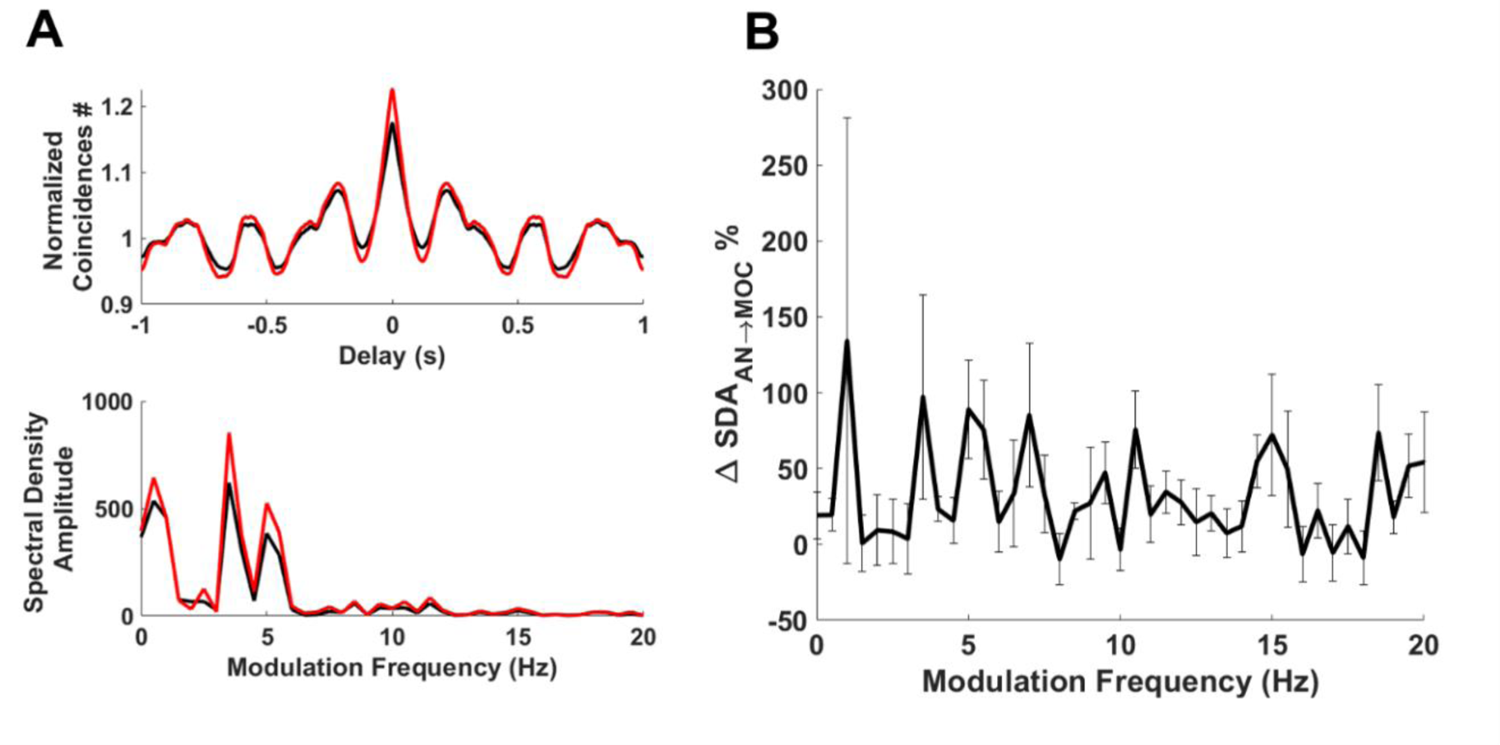
Efferent feedback improves envelope encoding for naturally-spoken sentences. **A**. Shuffled-Correlogram Sumcors (upper panel) were calculated for the naturally-spoken sentence [125], ‘the steady drip is worse than the drenching rain’ (s86; The MAVA corpus [126]), using low spontaneous rate auditory nerve fiber output between 1-2kHz with (*red line*) and without (*black line*) efferent feedback. A longer, 1 sec delay-window was used compared to the single word presentation; in addition, inverted triangular compensation was implemented to compensate for large delays relative to signal length (Rallapalli and Heinz 2016). The envelope power spectral density (lower panel) was computed both with (*red line*) and without (*black line*) efferent feedback by computing Fourier transforms of the above Sumcors with a <1Hz spectral resolution. Efferent feedback was conducive to larger envelope responses, especially at low modulation frequencies associated with words and syllables. **B**. Mean percentage change (±SEM) in power spectral density amplitude (ΔSDA_AN⇾MOC_) with efferent feedback included was computed using 6 sentences (s7, s26, s37, s42, s86, s164; The MAVA corpus) for modulation frequencies below 20 Hz. As for the single example in A, adding efferent feedback improved envelope encoding across most modulation frequencies.

**Figure supplement 3.**
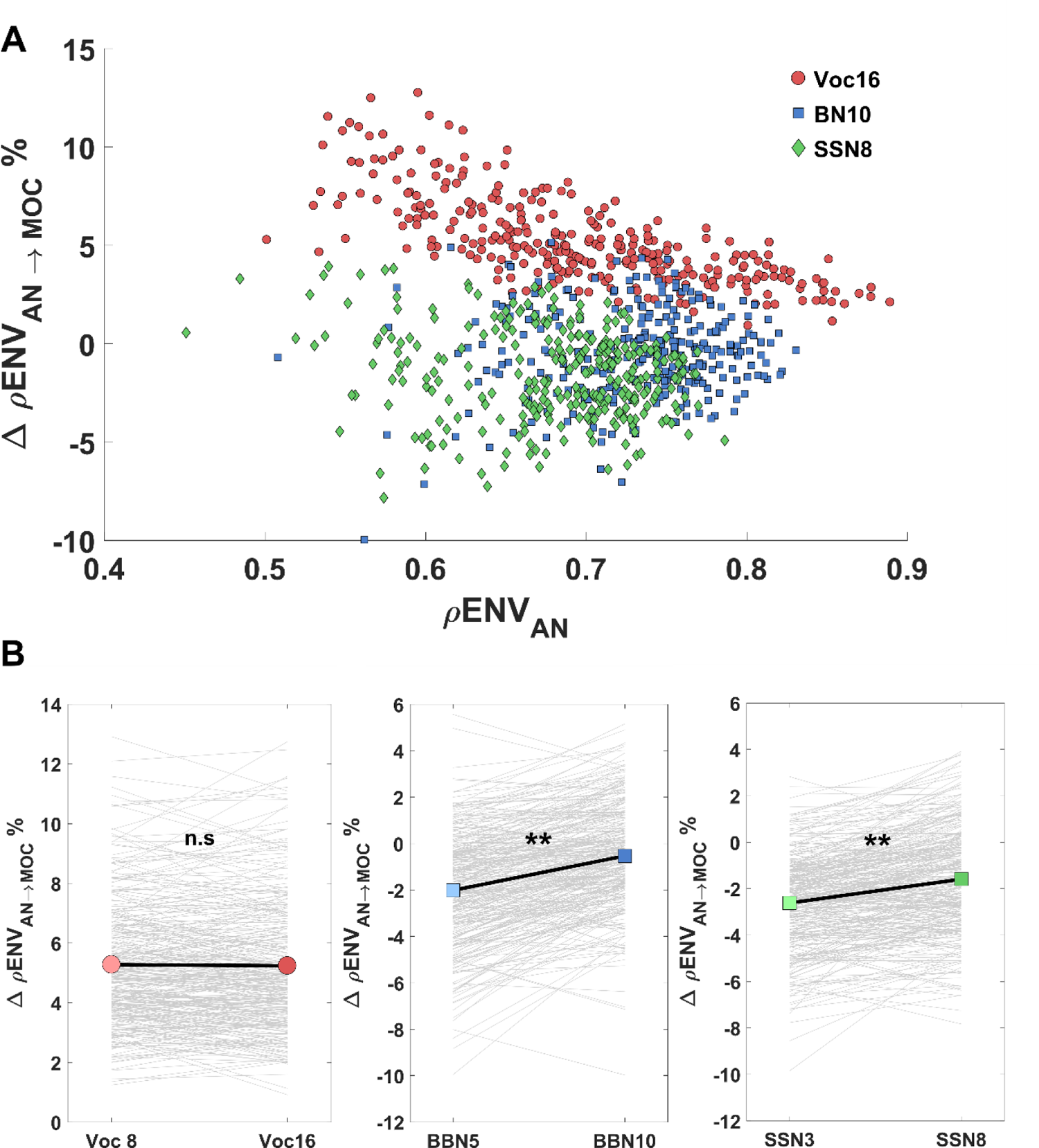
Presentation of degraded speech tokens with higher SNRs/more channels. **A.** Comparing ΔρENVs for 300 words after introduction of efferent feedback for Voc16, BBN10 and SSN8 stimuli. Mean percentage changes in ρENVs (calculated across 7 frequencies between 1-2kHz) after adding efferent feedback were plotted against ρENV without efferent feedback for degraded versions of 300 words (each symbol represents one word). ΔρENVs were always positive for Voc16 words (*pink circles*, D) (Max-Min ΔρENV for Voc8: +12.75 to +0.91%), appearing largest for words with lowest ρENVs without efferent feedback. This relationship was absent for BN5 (*light blue squares*, D) and SSN3 (*green diamonds*, D) words whose ΔρENV ranges spanned the baseline as they did for BN and SSN stimuli with lower SNRs (Max-Min ΔρENV for BN10: +5.16 to −9.97%; Max-Min ΔρENV for SSN8: +3.92 to −7.83%). **B.** Mean ΔρENV for BN10 and SSN8, however, increased significantly compared to their lower SNR counterparts, BN5 and SSN3 (mean ΔρENV for BN5 Vs BN10 = −2.01 ± 0.14 Vs. −0.5294 ± 0.13 [t(299) = −15.32, p *=* 0.0001]; mean ΔρENV for SSN3 Vs SSN8 = −2.62 ± 0.12 Vs. −1.59 ± 0.13, [t(299) = −10.89, p *=* 0.0001]). Mean ΔρENV for Voc16, on the other hand, was not significantly different from the Voc8 condition (mean ΔρENV for Voc8 Vs Voc 16= +5.30 ± 0.12% Vs. +5.24 ± 0.13%, [t(299) = 0.6535, p = 0.51]).

**Figure supplement 4.**
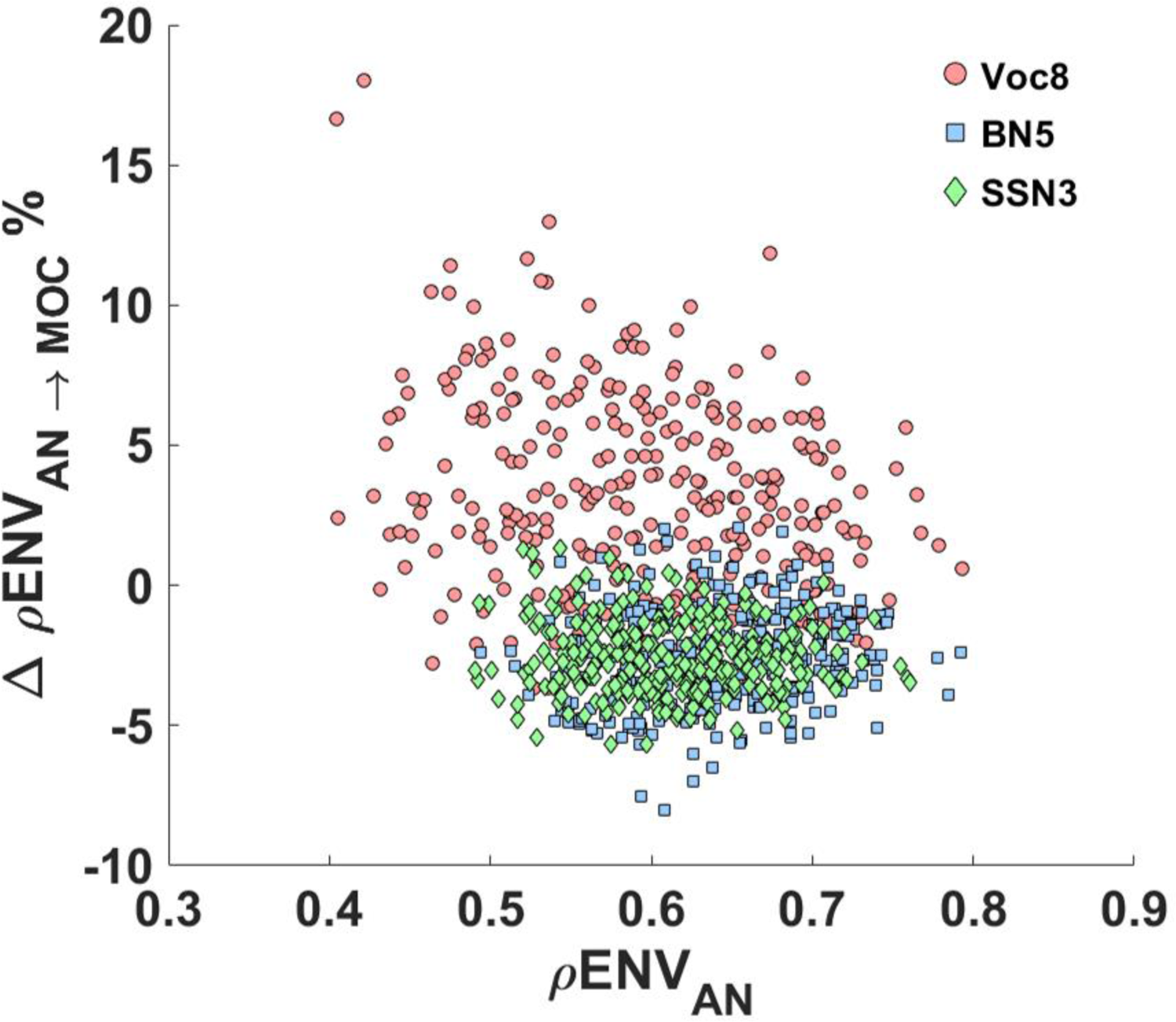
Percentage change in envelope encoding of high spontaneous rate (HSR) auditory nerve (AN) fibers after introduction of efferent feedback. ΔρENVs for 300 words (in their three degraded forms) were calculated as in Figure 2D; however, HSR AN fibre output was used here. Although ΔρENVs for Voc8 words (*pink circles*) varied greatly and included negative values (Max-Min ΔρENV for Voc8 = +16.87 to − 1.94%), mean ΔρENV was significantly positive (mean ΔρENV for Voc8 =+5.42 ± 0.18%, [t (299) = 30.17, p = 0.0001]). Note that values of ρENV_AN_ for Voc8 were significantly smaller here compared to matched values for low SR (LSR) AN fibers (mean ρENV_AN-HSR_ for Voc8 = 0.58 ± 0.01 *vs.* mean ρENV_AN-LSR_ for Voc8 = 0.70 ± 0.01, [t(299) = −39.21, p = 0.0001]). In contrast, the distributions of ΔρENVs for BN5 (*light blue squares*) and SSN3 (*green diamonds*) appeared more compact (Max-Min Range ΔρENV for BN5 words = +2.06 to −8.04%; Max-Min ΔρENV for SSN3 = +1.32 to −5.67 %); however, as for LSR AN fibre results, both mean ΔρENVs for HSR AN fibers were significantly negative overall (mean ΔρENV for BN5 = −2.54 ± 0.10, [t(299) = −24.92, p = 0.0001]; mean ΔρENV for SSN3 = −2.50 ± 0.08, [t(299) = −32.64, p = 0.0001]).

**Figure supplement 5.**
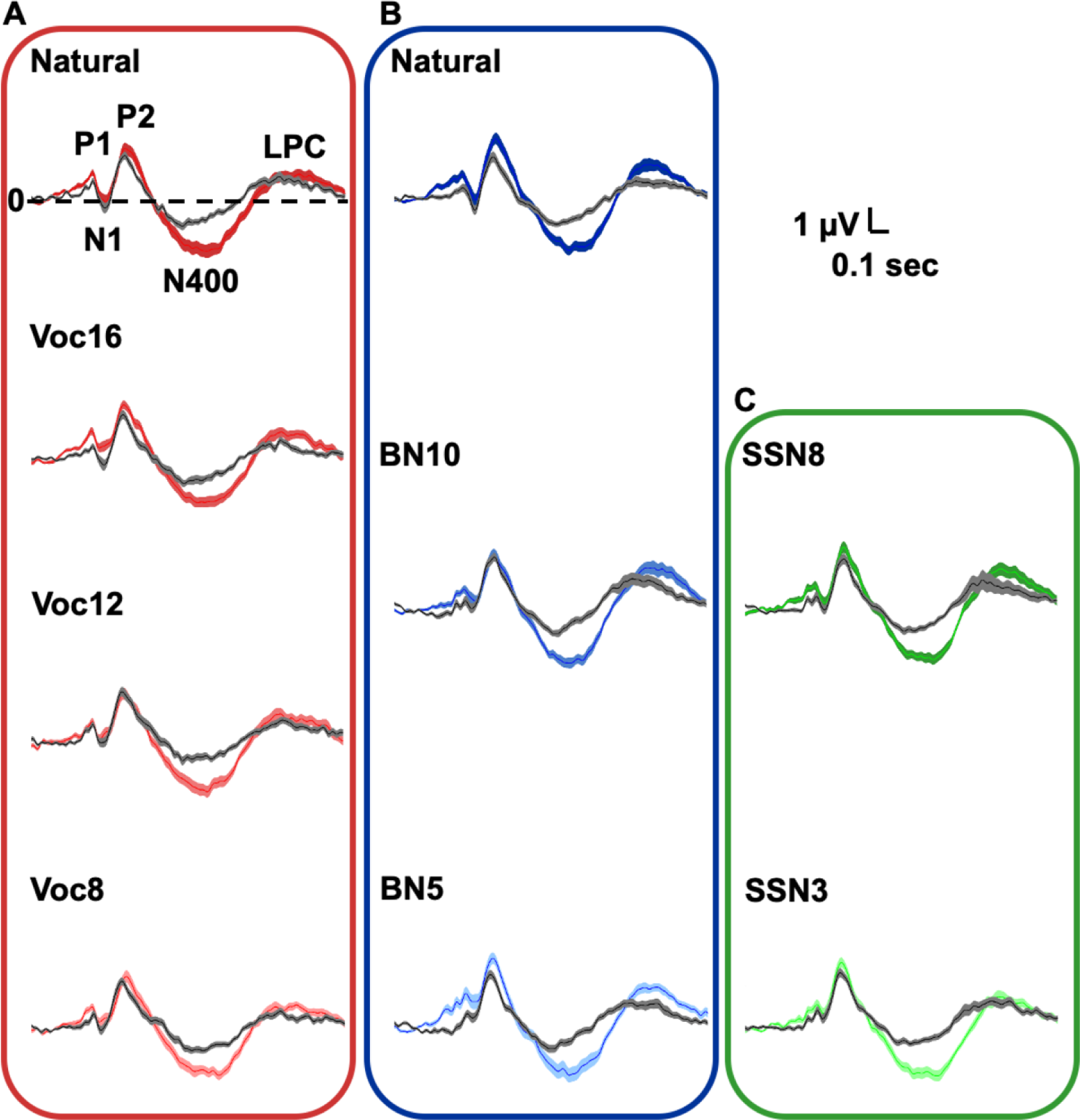
Cortical evoked potentials during active and passive speech perception. Thick lines and shaded areas represent means and standard errors of the mean respectively. Within conditions analysis showed that, for all speech manipulations, the magnitude of P1, P2 and N400 potentials were enhanced during active (*color lines)* when compared to the passive (*gray lines*) listening conditions, whilst N1 tended to be less negative in the active task. Late Positivity Complex (LPC) magnitude was only significantly enhanced during the active listening of speech in noise. **A.** ERP components in natural and all noise-vocoded manipulations: P1: [F (1,24) = 6.36, p = 0.02], N1: [F (1, 24) = 16.03, p = 0.001], P2: [F (1, 24) = 12.30, p = 0.002], N400: [F (1, 24) = 31.82, p = 0.0001], LPC: [F(1,24)=5.29, p = 0.03]. **B.** ERPs during natural (different population than noise-vocoded experiment) and all babble noise manipulations (n=29): P1: [F (1, 28) = 24.47, p = 0.0001], N1: [F (1, 28) = 10.46, p = 0.003], P2: [F (1, 28) = 10.65, p = 0.003], N400: [F (1, 28) = 62.16, p = 0.0001, LPC: [F(1,28)=10.55, p =0.003]. **C.** ERP components during speech-shaped noise manipulations (n=29): P1: [F (1, 28) = 22.98, p = 0.0001], N1: [F (1, 28) = 6.07, p = 0.02], P2: [F (1, 28) = 18.10, p = 0.001] and N400: [F (1, 28) = 60.75, p = 0.0001]), LPC: [F(1,28)=10.76, p =0.003].

**Figure supplement 6.**
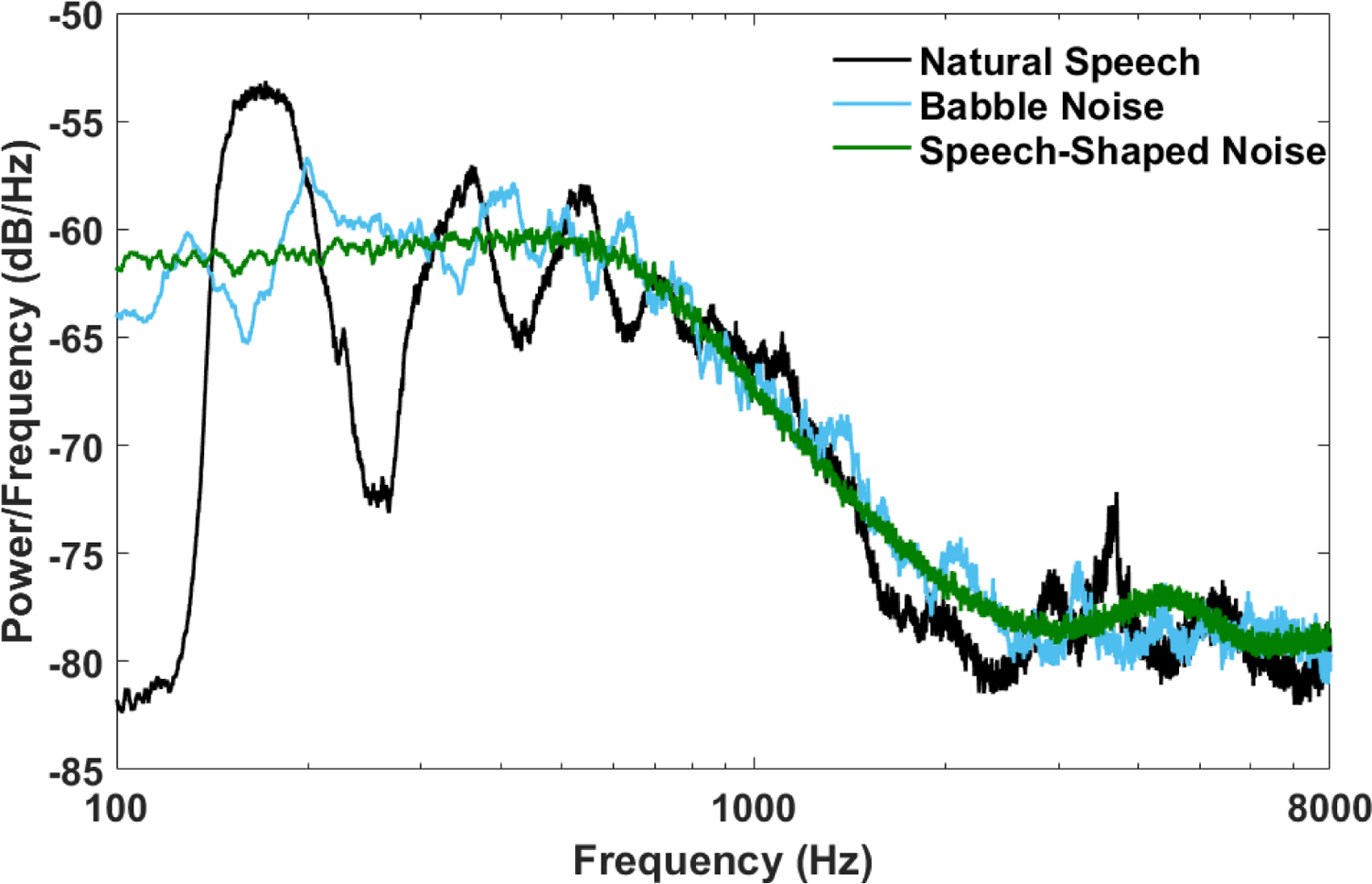
Comparison of Long-Term Average Spectra (LTAS) for natural speech, babble and speech-shaped noises. Power spectrum density estimates were calculated for 300 concatenated natural speech tokens (those presented to the MAP_BS model) and 60 seconds of 8-talker babble noise and speech-shaped noise; all acoustic stimuli had been normalised to 65 dB. The upper root-mean square envelopes, generated using 300-point sliding windows, are shown for the different conditions.

**Table supplement 1.** Hodgkin-Huxley model parameters and synaptic properties *LSR/HSR AN Fiber* = Low/High Spontaneous Rate Auditory Nerve Fiber; *LSR/HSR syn =* Low/High Spontaneous Inner Hair Cell Synapse; *VNTB* = Ventral Nucleus of the Trapezoid Body. ‘*ipsi*’ and ‘*contra*’ is relative to the simulated ear or cell in question. *CF* is the characteristic frequency of AN fiber tested and represents within-channel for input bandwidth. *LP dendritic filter* represents cut-off frequency of low-pass dendritic filter used to determine unitary synaptic conductance (*^g̅^_syn_*) at time, *t*, as follows:

|  | LSR AN Fiber | HSR AN Fiber | T Stellate | D Stellate | VNTB |
| --- | --- | --- | --- | --- | --- |
| Cell number | 200/CF/ear | 200/CF/ear | 50/CF/side | 50/CF/side | 50/CF/side |
| Cell type inputs: | ['ipsi' LSR syn] | ['ipsi' HR syn] | ['ipsi' AN (HSR); 'ipsi' VCN(chD)] | ['ipsi' AN(HSR)] | ['contra' VCN (ChT); 'ipsi' VCN (ChT)] |
| Input Bandwidth (Octaves) | [CF] | [CF] | [CF; CF] | [2] | [CF; CF] |
| Number of inputs | [1] | [1] | [15; 5] | [25] | [20; 5] |
| Max Synaptic Current (nA) | [30] | [40] | [2; 10] | [20] | [3; 3] |
| Synaptic Delay (ms) | [1.1] | [1.1] | [1.9; 1.9] | [1.9] | [1.9; 1.9] |
| LP dendritic filter (Hz) | [1000] | [1000] | [50; 50] | [500] | [70; 70] |
| Max Conductance (nS): |  |  |  |  |  |
| $\bar{g}_{Na}$ | [120] | [120] | [120] | [120] | [120] |
| $\bar{g}_{KLT}$ | [5] | [10] | [20] | [25] | [5] |
| $\bar{g}_{KHT}$ | [10] | [20] | [30] | [25] | [10] |
| $\bar{g}_H$ | [0] | [5] | [0] | [0] | [0] |
| $\bar{g}_L$ | [2.9] | [2.9] | [2.9] | [2.9] | [2.9] |
| Reversal Potentials (mV): |  |  |  |  |  |
| $E_{Na}$ | [+55] | [+55] | [+55] | [+55] | [+55] |
| $E_K$ | [-80] | [-80] | [-80] | [-80] | [-80] |
| $E_H$ | N/A | [-43] | N/A | N/A | N/A |
| $E_L$ | [-62] | [-61] | [-59] | [-63] | [-64] |
| Membrane Capacitance (pF): |  |  |  |  |  |
| $C_m$ | [1] | [1] | [1] | [1] | [1] |
| Excitatory/Inhibitory output | N/A | Excitation | Excitation | Inhibition | MOC attenuation |

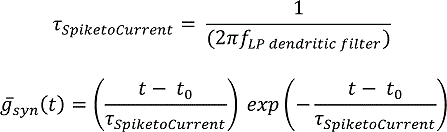

Where *t_0_* is the onset of the synaptic event. Current per input per spike was calculated by dividing *Max synaptic current* by *Number of inputs*. The above conductance and synaptic parameters are not absolute values in the physiological sense, instead mirroring key features of the physiology with a view to reproducing the time course of spiking activity and the qualitative differences of neural response patterns.

**Table supplement 2.**
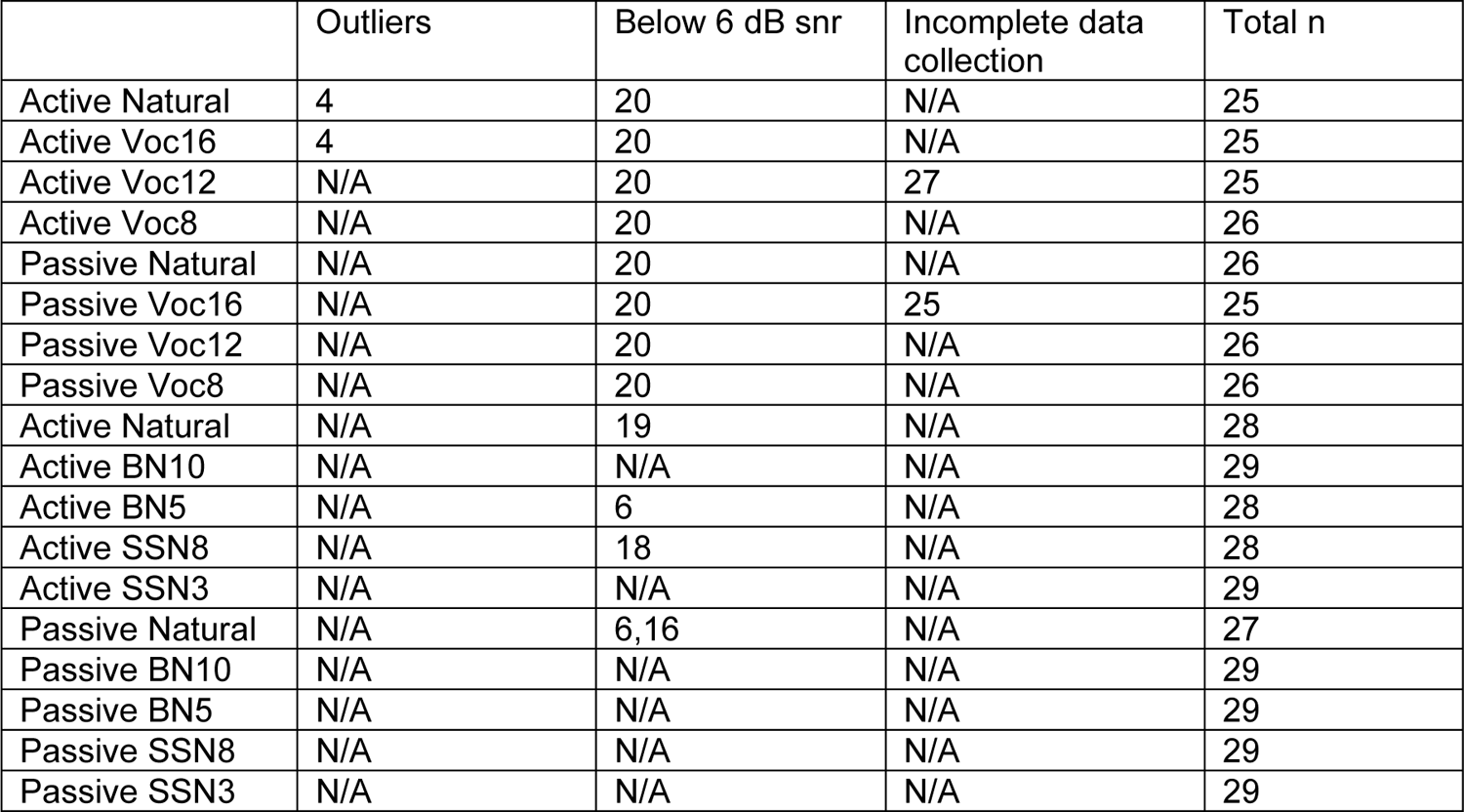
Subjects removed for CEOAEs inhibition analysis

**Table supplement 3.** Subjects removed for ABRs analysis

|  | Outliers | Below noise floor | Incomplete data collection | Total n |
| --- | --- | --- | --- | --- |
| Noise-vocoded-Wave V | 3 | 5 | 25,27 | 23 |
| Noise-vocoded-Wave III | 3 | 5, 12, 16 | 25,27 | 21 |
| Noise-vocoded-Ratio | 3 | 5, 12, 16 | 25,27 | 21 |
| Babble Noise-Wave V | 10 | 16 | N/A | 27 |
| Babble Noise-Wave III | 5, 27 | 6,16 | N/A | 24 |
| Babble Noise-Ratio | 5, 19 | 6,16 | N/A | 25 |
| Speech-Shaped noise-Wave V | 10 | 16 | N/A | 27 |
| Speech-Shaped noise-Wave III | 5, 27 | 6,13,16 | N/A | 25 |
| Speech-Shaped noise-Ratio | 5, 19 | 6,16 | N/A | 25 |

## Notes

### Competing Interest Statement

The authors have declared no competing interest.

## References

1. Lesica NA. Why do hearing aids fail to restore normal auditory perception? Trends Neurosci. 2018;41: 174–185.

2. Lehmann J, Christen N, Barilan YM, Gannot I. Age-related hearing loss, speech understanding and cognitive technologies. Int J Speech Technol. 2021. doi:10.1007/s10772-021-09817-z

3. Chien W, Lin FR. Prevalence of hearing aid use among older adults in the United States. Arch Intern Med. 2012;172: 292–293.

4. Simpson AN, Matthews LJ, Cassarly C, Dubno JR. Time From Hearing Aid Candidacy to Hearing Aid Adoption: A Longitudinal Cohort Study. Ear Hear. 2019;40: 468–476.

5. Irace AL, Sharma RK, Reed NS, Golub JS. Smartphone-Based Applications to Detect Hearing Loss: A Review of Current Technology. J Am Geriatr Soc. 2021;69: 307–316.

6. Darrow KN, Maison SF, Liberman MC. Cochlear efferent feedback balances interaural sensitivity. Nat Neurosci. 2006;9: 1474–1476.

7. Grothe B, Pecka M. The natural history of sound localization in mammals--a story of neuronal inhibition. Front Neural Circuits. 2014;8: 116.

8. Smith DW, Keil A. The biological role of the medial olivocochlear efferents in hearing: separating evolved function from exaptation. Front Syst Neurosci. 2015;9: 12.

9. Ahveninen J, Jääskeläinen IP, Raij T, Bonmassar G, Devore S, Hämäläinen M, et al. Task-modulated “what” and “where” pathways in human auditory cortex. Proc Natl Acad Sci U S A. 2006;103: 14608–14613.

10. Knudsen EI. Fundamental components of attention. Annu Rev Neurosci. 2007;30: 57–78.

11. Mulders WH, Robertson D. Evidence for direct cortical innervation of medial olivocochlear neurones in rats. Hear Res. 2000;144: 65–72.

12. Xiao Z, Suga N. Modulation of cochlear hair cells by the auditory cortex in the mustached bat. Nat Neurosci. 2002;5: 57–63.

13. Dragicevic CD, Aedo C, León A, Bowen M, Jara N, Terreros G, et al. The olivocochlear reflex strength and cochlear sensitivity are independently modulated by auditory cortex microstimulation. J Assoc Res Otolaryngol. 2015;16: 223–240.

14. Ashmore J, Avan P, Brownell WE, Dallos P, Dierkes K, Fettiplace R, et al. The remarkable cochlear amplifier. Hear Res. 2010;266: 1–17.

15. Almishaal A, Jennings SG. Effects of a precursor on amplitude modulation detection are consistent with efferent feedback. J Acoust Soc Am. 2016;139: 2155–2155.

16. Winslow RL, Sachs MB. Effect of electrical stimulation of the crossed olivocochlear bundle on auditory nerve response to tones in noise. J Neurophysiol. 1987;57: 1002–1021.

17. Guinan JJ Jr, Gifford ML. Effects of electrical stimulation of efferent olivocochlear neurons on cat auditory-nerve fibers. I. Rate-level functions. Hear Res. 1988;33: 97–113.

18. Faye-Lund H. Projection from the inferior colliculus to the superior olivary complex in the albino rat. Anat Embryol. 1986;175: 35–52.

19. Mulders WH, Robertson D. Effects on cochlear responses of activation of descending pathways from the inferior colliculus. Hear Res. 2000;149: 11–23.

20. Suthakar K, Ryugo DK. Descending projections from the inferior colliculus to medial olivocochlear efferents: Mice with normal hearing, early onset hearing loss, and congenital deafness. Hear Res. 2017;343: 34–49.

21. Guinan JJ Jr. Olivocochlear efferents: Their action, effects, measurement and uses, and the impact of the new conception of cochlear mechanical responses. Hear Res. 2018;362: 38–47.

22. de Boer J, Thornton ARD, Krumbholz K. What is the role of the medial olivocochlear system in speech-in-noise processing? J Neurophysiol. 2012;107: 1301–1312.

23. Mishra SK, Lutman ME. Top-down influences of the medial olivocochlear efferent system in speech perception in noise. PLoS One. 2014;9: e85756.

24. Mertes IB, Johnson KM, Dinger ZA. Olivocochlear efferent contributions to speech-in-noise recognition across signal-to-noise ratios. J Acoust Soc Am. 2019;145: 1529.

25. Kemp DT. Stimulated acoustic emissions from within the human auditory system. J Acoust Soc Am. 1978;64: 1386–1391.

26. Collet L, Kemp DT, Veuillet E, Duclaux R, Moulin A, Morgon A. Effect of contralateral auditory stimuli on active cochlear micro-mechanical properties in human subjects. Hear Res. 1990;43: 251–261.

27. Giraud AL, Garnier S, Micheyl C, Lina G, Chays A, Chéry-Croze S. Auditory efferents involved in speech-in-noise intelligibility. Neuroreport. 1997;8: 1779–1783.

28. Harkrider AW, Bowers CD. Evidence for a cortically mediated release from inhibition in the human cochlea. J Am Acad Audiol. 2009;20: 208–215.

29. Wagner W, Frey K, Heppelmann G, Plontke SK, Zenner H-P. Speech-in-noise intelligibility does not correlate with efferent olivocochlear reflex in humans with normal hearing. Acta Otolaryngol. 2008;128: 53–60.

30. Stuart A, Butler AK. Contralateral suppression of transient otoacoustic emissions and sentence recognition in noise in young adults. J Am Acad Audiol. 2012;23: 686–696.

31. Schaette R, McAlpine D. Tinnitus with a normal audiogram: physiological evidence for hidden hearing loss and computational model. J Neurosci. 2011;31: 13452–13457.

32. Kahneman D. Attention and effort. Prentice-Hall; 1973.

33. Meddis R. MAP-BS a Matlab Auditory Processing software platform for studying Auditory BrainStem activity. 2016 [cited 3 Jun 2020]. doi:10.13140/RG.2.2.30627.45603

34. Costalupes JA, Young ED, Gibson DJ. Effects of continuous noise backgrounds on rate response of auditory nerve fibers in cat. J Neurophysiol. 1984;51: 1326–1344.

35. Kujawa SG, Liberman MC. Synaptopathy in the noise-exposed and aging cochlea: Primary neural degeneration in acquired sensorineural hearing loss. Hear Res. 2015;330: 191–199.

36. Huet A, Desmadryl G, Justal T, Nouvian R, Puel J-L, Bourien J. The Interplay Between Spike-Time and Spike-Rate Modes in the Auditory Nerve Encodes Tone-In-Noise Threshold. J Neurosci. 2018;38: 5727–5738.

37. Heinz MG, Swaminathan J. Quantifying envelope and fine-structure coding in auditory nerve responses to chimaeric speech. J Assoc Res Otolaryngol. 2009;10: 407–423.

38. Liberman MC. Auditory-nerve response from cats raised in a low-noise chamber. J Acoust Soc Am. 1978;63: 442–455.

39. Winter IM, Palmer AR. Intensity coding in low-frequency auditory-nerve fibers of the guinea pig. J Acoust Soc Am. 1991;90: 1958–1967.

40. Heinz MG, Young ED. Response growth with sound level in auditory-nerve fibers after noise-induced hearing loss. J Neurophysiol. 2004;91: 784–795.

41. Carney LH. Supra-Threshold Hearing and Fluctuation Profiles: Implications for Sensorineural and Hidden Hearing Loss. J Assoc Res Otolaryngol. 2018. doi:10.1007/s10162-018-0669-5

42. Encina-Llamas G, Harte JM, Dau T, Shinn-Cunningham B, Epp B. Investigating the effect of cochlear synaptopathy on envelope following responses using a model of the auditory nerve. J Assoc Res Otolaryngol. 2019;20: 363–382.

43. Pujol R. Cerveau auditif. [cited 10 Mar 2021]. Available: http://www.cochlea.eu/cerveau-auditif

44. Getzmann S, Falkenstein M, Wascher E. ERP correlates of auditory goal-directed behavior of younger and older adults in a dynamic speech perception task. Behav Brain Res. 2015;278: 435–445.

45. Grunwald T, Boutros NN, Pezer N, von Oertzen J, Fernández G, Schaller C, et al. Neuronal substrates of sensory gating within the human brain. Biol Psychiatry. 2003;53: 511–519.

46. Potts GF. An ERP index of task relevance evaluation of visual stimuli. Brain Cogn. 2004;56: 5–13.

47. Juottonen K, Revonsuo A, Lang H. Dissimilar age influences on two ERP waveforms (LPC and N400) reflecting semantic context effect. Brain Res Cogn Brain Res. 1996;4: 99–107.

48. Stuss DT, Picton TW, Cerri AM, Leech EE, Stethem LL. Perceptual closure and object identification: electrophysiological responses to incomplete pictures. Brain Cogn. 1992;19: 253–266.

49. Francis AL, MacPherson MK, Chandrasekaran B, Alvar AM. Autonomic Nervous System Responses During Perception of Masked Speech may Reflect Constructs other than Subjective Listening Effort. Front Psychol. 2016;7: 263.

50. Pichora-Fuller MK, Kramer SE, Eckert MA, Edwards B, Hornsby BWY, Humes LE, et al. Hearing Impairment and Cognitive Energy: The Framework for Understanding Effortful Listening (FUEL). Ear Hear. 2016;37 Suppl 1: 5S–27S.

51. Kutas M, Federmeier KD. Thirty years and counting: finding meaning in the N400 component of the event-related brain potential (ERP). Annu Rev Psychol. 2011;62: 621–647.

52. Puel JL, Bonfils P, Pujol R. Selective attention modifies the active micromechanical properties of the cochlea. Brain Res. 1988;447: 380–383.

53. Wittekindt A, Kaiser J, Abel C. Attentional modulation of the inner ear: a combined otoacoustic emission and EEG study. J Neurosci. 2014;34: 9995–10002.

54. Bowen M, Terreros G, Moreno-Gómez FN, Ipinza M, Vicencio S, Robles L, et al. The olivocochlear reflex strength in awake chinchillas is relevant for behavioural performance during visual selective attention with auditory distractors. Sci Rep. 2020;10: 14894.

55. Kawase T, Delgutte B, Liberman MC. Antimasking effects of the olivocochlear reflex. II. Enhancement of auditory-nerve response to masked tones. J Neurophysiol. 1993;70: 2533– 2549.

56. Marian V, Lam TQ, Hayakawa S, Dhar S. Spontaneous Otoacoustic Emissions Reveal an Efficient Auditory Efferent Network. J Speech Lang Hear Res. 2018;61: 2827–2832.

57. Saiz-Alía M, Miller P, Reichenbach T. Otoacoustic emissions evoked by the time-varying harmonic structure of speech. eNeuro. 2021. doi:10.1523/ENEURO.0428-20.2021

58. Lavie N. Distracted and confused?: selective attention under load. Trends Cogn Sci. 2005;9: 75–82.

59. Lilaonitkul W, Guinan JJ Jr. Human medial olivocochlear reflex: effects as functions of contralateral, ipsilateral, and bilateral elicitor bandwidths. J Assoc Res Otolaryngol. 2009;10: 459–470.

60. Boothalingam S, Purcell D, Scollie S. Influence of 100Hz amplitude modulation on the human medial olivocochlear reflex. Neurosci Lett. 2014;580: 56–61.

61. Kalaiah MK, Nanchirakal JF, Kharmawphlang L, Noronah SC. Contralateral suppression of transient evoked otoacoustic emissions for various noise signals. Hearing, Balance and Communication. 2017;15: 84–90.

62. Lopez-Poveda EA, Eustaquio-Martín A, Stohl JS, Wolford RD, Schatzer R, Wilson BS. A Binaural Cochlear Implant Sound Coding Strategy Inspired by the Contralateral Medial Olivocochlear Reflex. Ear Hear. 2016;37: e138–48.

63. Terreros G, Delano PH. Corticofugal modulation of peripheral auditory responses. Front Syst Neurosci. 2015;9: 134.

64. Hausfeld L, Shiell M, Formisano E, Riecke L. Cortical processing of distracting speech in noisy auditory scenes depends on perceptual demand. Neuroimage. 2021;228: 117670.

65. Heggdal POL, Aarstad HJ, Brännström J, Vassbotn FS, Specht K. An fMRI-study on single-sided deafness: Spectral-temporal properties and side of stimulation modulates hemispheric dominance. Neuroimage Clin. 2019;24: 101969.

66. Khalfa S, Bougeard R, Morand N, Veuillet E, Isnard J, Guenot M, et al. Evidence of peripheral auditory activity modulation by the auditory cortex in humans. Neuroscience. 2001;104: 347– 358.

67. Maison SF, Liberman MC. Predicting vulnerability to acoustic injury with a noninvasive assay of olivocochlear reflex strength. J Neurosci. 2000;20: 4701–4707.

68. Taranda J, Maison SF, Ballestero JA, Katz E, Savino J, Vetter DE, et al. A point mutation in the hair cell nicotinic cholinergic receptor prolongs cochlear inhibition and enhances noise protection. PLoS Biol. 2009;7: e18.

69. Boero LE, Castagna VC, Terreros G, Moglie MJ, Silva S, Maass JC, et al. Preventing presbycusis in mice with enhanced medial olivocochlear feedback. Proc Natl Acad Sci U S A. 2020;117: 11811–11819.

70. Wiederhold ML, Kiang NY. Effects of electric stimulation of the crossed olivocochlear bundle on single auditory-nerve fibers in the cat. J Acoust Soc Am. 1970;48: 950–965.

71. Cooper NP, Guinan JJ Jr. Efferent-mediated control of basilar membrane motion. J Physiol. 2006;576: 49–54.

72. Murugasu E, Russell IJ. The effect of efferent stimulation on basilar membrane displacement in the basal turn of the guinea pig cochlea. J Neurosci. 1996;16: 325–332.

73. Marrufo-Pérez MI, Eustaquio-Martín A, Lopez-Poveda EA. Adaptation to Noise in Human Speech Recognition Unrelated to the Medial Olivocochlear Reflex. J Neurosci. 2018;38: 4138– 4145.

74. Brown GJ, Ferry RT, Meddis R. A computer model of auditory efferent suppression: implications for the recognition of speech in noise. J Acoust Soc Am. 2010;127: 943–954.

75. Clark NR, Brown GJ, Jürgens T, Meddis R. A frequency-selective feedback model of auditory efferent suppression and its implications for the recognition of speech in noise. J Acoust Soc Am. 2012;132: 1535–1541.

76. Yasin I, Drga V, Liu F, Demosthenous A, Meddis R. Optimizing Speech Recognition Using a Computational Model of Human Hearing: Effect of Noise Type and Efferent Time Constants. IEEE Access. 2020;8: 56711–56719.

77. Messing DP, Delhorne L, Bruckert E, Braida LD, Ghitza O. A non-linear efferent-inspired model of the auditory system; matching human confusions in stationary noise. Speech Commun. 2009;51: 668–683.

78. Mertes IB, Wilbanks EC, Leek MR. Olivocochlear Efferent Activity Is Associated With the Slope of the Psychometric Function of Speech Recognition in Noise. Ear Hear. 2018;39: 583–593.

79. Marrufo-Pérez MI, Sturla-Carreto DDP, Eustaquio-Martín A, Lopez-Poveda EA. Adaptation to Noise in Human Speech Recognition Depends on Noise-Level Statistics and Fast Dynamic-Range Compression. J Neurosci. 2020;40: 6613–6623.

80. Wojtczak M, Klang AM, Torunsky NT. Exploring the Role of Medial Olivocochlear Efferents on the Detection of Amplitude Modulation for Tones Presented in Noise. J Assoc Res Otolaryngol. 2019;20: 395–413.

81. Miller GA, Licklider JCR. The intelligibility of interrupted speech. J Acoust Soc Am. 1950;22: 167–173.

82. Cooke M. A glimpsing model of speech perception in noise. J Acoust Soc Am. 2006;119: 1562–1573.

83. Li N, Loizou PC. Factors influencing glimpsing of speech in noise. J Acoust Soc Am. 2007;122: 1165–1172.

84. Jørgensen S, Ewert SD, Dau T. A multi-resolution envelope-power based model for speech intelligibility. J Acoust Soc Am. 2013;134: 436–446.

85. Relaño-Iborra H, May T, Zaar J, Scheidiger C, Dau T. Predicting speech intelligibility based on a correlation metric in the envelope power spectrum domain. J Acoust Soc Am. 2016;140: 2670.

86. de Boer J, Thornton ARD. Neural correlates of perceptual learning in the auditory brainstem: efferent activity predicts and reflects improvement at a speech-in-noise discrimination task. J Neurosci. 2008;28: 4929–4937.

87. Ding N, Simon JZ. Adaptive temporal encoding leads to a background-insensitive cortical representation of speech. J Neurosci. 2013;33: 5728–5735.

88. Vanthornhout J, Decruy L, Wouters J, Simon JZ, Francart T. Speech Intelligibility Predicted from Neural Entrainment of the Speech Envelope. J Assoc Res Otolaryngol. 2018;19: 181– 191.

89. Brodbeck C, Jiao A, Hong LE, Simon JZ. Neural speech restoration at the cocktail party: Auditory cortex recovers masked speech of both attended and ignored speakers. PLoS Biol. 2020;18: e3000883.

90. Zeng F-G, Nie K, Liu S, Stickney G, Del Rio E, Kong Y-Y, et al. On the dichotomy in auditory perception between temporal envelope and fine structure cues. J Acoust Soc Am. 2004;116: 1351–1354.

91. Lorenzi C, Gilbert G, Carn H, Garnier S, Moore BCJ. Speech perception problems of the hearing impaired reflect inability to use temporal fine structure. Proc Natl Acad Sci U S A. 2006;103: 18866–18869.

92. Shamma S, Lorenzi C. On the balance of envelope and temporal fine structure in the encoding of speech in the early auditory system. J Acoust Soc Am. 2013;133: 2818–2833.

93. Ding N, Chatterjee M, Simon JZ. Robust cortical entrainment to the speech envelope relies on the spectro-temporal fine structure. Neuroimage. 2014;88: 41–46.

94. Viswanathan V, Bharadwaj HM, Shinn-Cunningham B, Heinz MG. Evaluating human neural envelope coding as the basis of speech intelligibility in noise. J Acoust Soc Am. 2019;145: 1717–1717.

95. Brown MC. Single-unit labeling of medial olivocochlear neurons: the cochlear frequency map for efferent axons. J Neurophysiol. 2014;111: 2177–2186.

96. Liberman LD, Liberman MC. Cochlear Efferent Innervation Is Sparse in Humans and Decreases with Age. J Neurosci. 2019;39: 9560–9569.

97. Forte AE, Etard O, Reichenbach T. The human auditory brainstem response to running speech reveals a subcortical mechanism for selective attention. Elife. 2017;6: e27203.

98. Brix R. The influence of attention on the auditory brain stem evoked responses. Preliminary report. Acta Otolaryngol. 1984;98: 89–92.

99. Lukas JH. Human auditory attention: the olivocochlear bundle may function as a peripheral filter. Psychophysiology. 1980;17: 444–452.

100. Hillyard SA, Hink RF, Schwent VL, Picton TW. Electrical signs of selective attention in the human brain. Science. 1973;182: 177–180.

101. Fujiwara N, Nagamine T, Imai M, Tanaka T, Shibasaki H. Role of the primary auditory cortex in auditory selective attention studied by whole-head neuromagnetometer. Brain Res Cogn Brain Res. 1998;7: 99–109.

102. Mesgarani N, David SV, Fritz JB, Shamma SA. Mechanisms of noise robust representation of speech in primary auditory cortex. Proc Natl Acad Sci U S A. 2014;111: 6792–6797.

103. Rabinowitz NC, Willmore BDB, King AJ, Schnupp JWH. Constructing noise-invariant representations of sound in the auditory pathway. PLoS Biol. 2013;11: e1001710.

104. Kell AJE, McDermott JH. Invariance to background noise as a signature of non-primary auditory cortex. Nat Commun. 2019;10. doi:10.1038/s41467-019-11710-y

105. Robinson BL, Harper NS, McAlpine D. Meta-adaptation in the auditory midbrain under cortical influence. Nat Commun. 2016;7: 13442.

106. Shaheen LA, Slee SJ, David SV. Task Engagement Improves Neural Discriminability in the Auditory Midbrain of the Marmoset Monkey. J Neurosci. 2021;41: 284–297.

107. Asokan MM, Williamson RS, Hancock KE, Polley DB. Sensory overamplification in layer 5 auditory corticofugal projection neurons following cochlear nerve synaptic damage. Nat Commun. 2018;9: 2468.

108. Lehiste I, Peterson GE. Linguistic Considerations in the Study of Speech Intelligibility. J Acoust Soc Am. 1959;31: 280–286.

109. Rastle K, Harrington J, Coltheart M. 358,534 nonwords: the ARC Nonword Database. Q J Exp Psychol A. 2002;55: 1339–1362.

110. Ritzer G, editor. Experimental Design. The Blackwell Encyclopedia of Sociology. Oxford, UK: John Wiley & Sons, Ltd; 2007.

111. Shannon RV, Zeng FG, Kamath V, Wygonski J, Ekelid M. Speech recognition with primarily temporal cues. Science. 1995;270: 303–304.

112. Keidser G, Ching T, Dillon H, Agung K, Brew C, Brewer S, et al. The National Acoustic Laboratories’ (NAL) CDs of Speech and Noise for Hearing Aid Evaluation: Normative Data and Potential Applications. Australian and New Zealand Journal of Audiology. 2002; 24: 16–35.

113. Boothalingam S, Purcell DW. Influence of the stimulus presentation rate on medial olivocochlear system assays. J Acoust Soc Am. 2015;137: 724–732.

114. Souza NN, Dhar S, Neely ST, Siegel JH. Comparison of nine methods to estimate ear-canal stimulus levels. J Acoust Soc Am. 2014;136: 1768–1787.

115. Hood LJ, Berlin CI, Hurley A, Cecola RP, Bell B. Contralateral suppression of transient-evoked otoacoustic emissions in humans: intensity effects. Hear Res. 1996;101: 113–118.

116. Murata K, Ito S, Horikawa J, Minami S. The acoustic middle ear muscle reflex in albino rats. Hear Res. 1986;23: 169–183.

117. Liberman MC, Guinan JJ Jr. Feedback control of the auditory periphery: anti-masking effects of middle ear muscles vs. olivocochlear efferents. J Commun Disord. 1998;31: 471– 82; quiz 483; 553.

118. Lee DJ, de Venecia RK, Guinan JJJr, Brown MC. Central auditory pathways mediating the rat middle ear muscle reflexes. Anat Rec A Discov Mol Cell Evol Biol. 2006;288: 358–369.

119. Oostenveld R, Fries P, Maris E, Schoffelen J-M. FieldTrip: Open source software for advanced analysis of MEG, EEG, and invasive electrophysiological data. Comput Intell Neurosci. 2011;2011: 156869.

120. Woodman GF. A brief introduction to the use of event-related potentials in studies of perception and attention. Atten Percept Psychophys. 2010;72: 2031–2046.

121. Don M, Elberling C. Evaluating residual background noise in human auditory brain-stem responses. J Acoust Soc Am. 1994;96: 2746–2757.

122. Faul F, Erdfelder E, Lang A-G, Buchner A. G*Power 3: a flexible statistical power analysis program for the social, behavioral, and biomedical sciences. Behav Res Methods. 2007;39: 175–191.

123. Liberman MC. Response properties of cochlear efferent neurons: monaural vs. binaural stimulation and the effects of noise. J Neurophysiol. 1988;60: 1779–1798.

124. Joris PX, Louage DH, Cardoen L, van der Heijden M. Correlation index: a new metric to quantify temporal coding. Hear Res. 2006;216–217: 19–30.

125. Rallapalli VH, Heinz MG. Neural Spike-Train Analyses of the Speech-Based Envelope Power Spectrum Model: Application to Predicting Individual Differences with Sensorineural Hearing Loss. Trends in Hearing. 2016;20: 2331216516667319.

126. Aubanel, V., Davis, C. Kim, J. The MAVA corpus. 2017. doi:10.4227/139/59a4c21a896a3

